# Characterization of variants associated with Cerebral Small Vessel Disease identifies a functional SNV in Versican

**DOI:** 10.64898/2026.03.16.712010

**Authors:** Jae-Ryeon Ryu, Ankita Narang, Quentin LeGrand, David-Alexandre Trégouët, Stephanie Debette, Sarah J. Childs

**Author notes:** **Correspondence:** Sarah Childs.

## Abstract

**Background:** Single nucleotide variants in the non-coding genome can significantly alter disease risk, but identifying the function of SNVs is a challenge. Increasing numbers of variants of unknown significance have been associated with the risk of Stroke, Cerebral Small Vessel Disease and burden of White Matter Hyperintensities, but without biological validation, the significance of these findings is uncertain.

**Methods:** We use a Multiplexed Parallel Reporter Assay (MPRA) to assay the function of 44 SNVs associated with Cerebral Small Vessel Disease and stroke in *EFEMP1, FOXF2, HAAO, KCNK3, NMT1, OPA1, RASL12, STAT3 and VCAN*. A functional SNV in versican was further probed using transcriptional reporter and Chromatin Immunoprecipitation assays.

**Results:** We identify 26 allele-specific enhancers in where the SNV either increases or decreases transcriptional activity. We validate rs13176921 as an SNV that significantly modulates an intronic enhancer in the matrisome protein Versican and show it affects mRNA levels of the gene. We identify that the transcription factor NKX3.1 binds to the region of this SNV.

**Conclusions:** Thus, using MRPA we were able to gain mechanistic insight into which GWAS-identified SNVs modulate transcriptional activity, and validate a functionally important SNV in Versican and provide a mechanism by which it controls its transcription.

## Introduction

Small vessels (arterioles, capillaries, and venules) provide blood flow to support cognitive tasks through an organized distribution system. The smallest vessels with the slowest flow are critical for delivery of oxygen and nutrients to tissues. For this reason, damage to small vessels can lead to significant tissue consequences. Cerebral small vessel disease (SVD) results in poor brain blood flow and leads to ~20% of stroke and 45% of vascular dementias ^1^. SVD originates from blood vessels and secondarily affects neurons. The global lifetime risk of ischemic stroke in humans is 18.3% ^2^, and ~25% of ischemic stroke is due to SVD. SVD is thought to be a lifespan disease that begins during development and worsens across the lifespan ^3^.

Brain blood vessels are critical for maintaining the blood-brain barrier (BBB) through establishment of tight and adherens junctions between endothelial cells. A tight endothelium requires direct contact with vascular mural cells (vascular smooth muscle cells (vSMCs) and pericytes)^4^, providing physical support, extracellular matrix (ECM) deposition and contractility^5^. Maturation of blood vessels requires coordination of reciprocal signalling among endothelium, mural cells, and ECM. In blood vessels, the majority of ECM is secreted by vascular mural cells ^6^ with the developmental period being of critical importance for its deposition. Mechanistically, many SVD-associated genes are members of the matrisome ^6^, the group of proteins that comprise or modify ECM.

SVD incidence becomes more common with aging, and genomic investigations, including genome-wide association studies (GWAS), have identified both monogenic ^7,8^, and common genetic variants that are associated with higher risk of the disease ^3,9,10^. In recent years, GWAS studies identified single nucleotide variants (SNVs) in known and novel genes associated with MRI-markers of cSVD, such as white matter hyperintensity volume, the most common neuroimaging feature of SVD ^3,9,11,12^. Using novel biophysical diffusion MRI techniques (neurite orientation and dispersion and density imaging (NODDI)) to identify white matter microstructure differences related to SVD in young adults, additional genetic associations emerged^13^. The role of these SNVs in modulating gene expression is unknown. Most SVD-associated SNVs are in either intergenic or intragenic regions in predicted cis-candidate regulatory elements (CCRE) enhancers on ENCODE^14^. Further, the role of many of the genes in vascular stabilization that map to the identified SNVs is unknown. Thus, both the variants themselves and many of the genes they are located near, are currently of unknown significance with respect to the mechanism of SVD.

In this study, we hypothesize that SVD associated variants are located in transcriptional enhancers. Transcriptional enhancers are located in intragenic and intergenic regions of the non-coding genome. While enhancers can be bioinformatically identified using epigenetic signatures and transcription factor (TF) binding, the precise mechanisms by which they regulate gene expression, the specific residues required for TF binding, and the target gene(s) they control must be determined through functional experiments. Thus, while GWA studies are an unbiased window into identifying disease-associated residues in the genome associated with disease, their utility is limited.

Multiplexed Reporter Assays (MPRAs) are unbiased assay of functional enhancer activity, testing the activity of alternative SNVs in a cell line-based assay ^15^. Using an MPRA, we assess the functional impact on enhancer activity of variants located near a selection of stroke and CSVD associated loci: *EFEMP1*, *FOXF2, HAAO, KCNK3*, *NMT1, OPA1, RASL12, STAT3* and *VCAN* ^3,9,11,13,16,17^. EFEMP1 (Fibulin 3) and *VCAN* (Versican), are both members of the human matrisome ^18^ and have strong expression in the aorta/artery^19^. *RASL12* is a very poorly studied gene, but, according to GTEx portal, it is expressed in a similar pattern to *EFEMP1* and *VCAN*. *HAAO, NMT1*, *OPA1*, *STAT3* and *KCNK3* are more broadly expressed, and are not part of the matrisome. Their connection to SVD and stroke likely occurs via a mechanism other than by ECM regulation.

Using MPRA, we identify allele-specific enhancers. For validation, we focus on intronic variants in Versican (*VCAN*) and identify rs13176921 as a critical variant. We show that the minor allele disturbs a conserved residue of the NKX3.1 binding site and show that NKX3.1 can be Chromatin immunoprecipitated (ChIP) to this site suggesting that rs13176921 may promote *VCAN* gene expression via NKX3.1 binding.

## Materials and methods

Primer sequences and reagent details are found in Supplemental Tables 1 and 2.

### MPRA library oligonucleotide synthesis

We designed a pool of 968 oligonucleotides from CSVD and stroke associated genes, containing 102 bp of flanking sequence upstream of the variant, the variant (major and minor variants), 102 bp of sequence downstream of the variant, a multi-enzyme cloning site, a short linker, a 10 bp barcode and a reverse adaptor (Supp. Table 3). An oligonucleotide pool was synthesized by Twist Bioscience (South San Francisco, CA). To amplify the library, 12 independent 50 µl PCR reactions were set up containing 1 ng the synthesized MPRA oligos, 10 µl of 5X Phusion HF buffer, 200 µM dNTPs, 1 unit Phusion HF DNA polymerase (New England Biolabs), 0.5uM MPRA oligo-f and MPRA oligo-r (Supp. Table 1) with the cycling conditions: 98 C for 30 sec, 15 cycles of (98 C for 10 sec, 63 C for 10 sec, 72 C for 15 sec), 72 C for 1min. The PCR-amplified MPRA oligos were pooled and purified with NucleoSpin Gel & PCR clean-up column (Macherey-Nagel).

### MPRA oligo library assembly

10 µg of pMPRAv3:Δluc:ΔxbaI (Addgene) was digested with SfiI (NEB) at 50C overnight and a single band extracted (NucleoSpin Gel & PCR clean-up column, Macherey-Nagel). To create the MPRA oligo library, 1.1 µg of MPRA oligos and 1 µg of digested pMPRAv3:Δluc:Δxbal were assembled using NEBuilder HiFi DNA Assembly (NEB) at 50C for 1 h. The reaction was purified with NucleoSpin PCR clean-up column and eluted in 20 µl elution buffer. 100-150 ng of the assembled MPRA oligo library was transformed into 50 µl of NEB 10-Beta high efficiency competent cells (NEB). 1 ml of NEB 10-Beta/Stable Outgrowth Medium (NEB) was added, and the cells were incubated at 37 C for 1 hour. To amplify the library, the recovered bacterial cells were cultured in four x 50 ml of LB with Ampicillin at 37C for 6 hours. After outgrowth cultures were pooled and the plasmid DNAs purified (NucleoBond Xtra Midi EF, (Macherey-Nagel).

### MPRA oligo:GFP library assembly

10 µg of MPRA oligo library was digested with AsiSI (NEB) at 37C overnight and a single band was gel-extracted (NucleoSpin Gel & PCR clean-up). A MiniGFP amplicon containing a minimal promoter, GFP ORF and a partial 3’ UTR (pMPRAv3:minP-GFP, Addgene) was PCR amplified with MiniP-GFP-f and MiniP-GFP-r primers ^20^ (Supp. Table 1). The MPRA library incorporating GFP library was created using 2 µg of MiniGFP amplicon and 1µg of AsiSI linearized MPRA oligo library and assembled using NEBuilder HiFi DNA Assembly at 50C for 90 min, then purified with NucleoSpin PCR clean-up column, and eluted in 20 µl elution buffer. The total elute was digested a second time to remove remaining uncut vectors by incubation with 25 U AsiSI, and 5 U RecBCD Plasmidsafe nuclease (NEB), 10µg BSA, 1 mM ATP in a 100 µl reaction at 37 C for overnight followed by NucleoSpin PCR clean-up column and eluted with 40 µl of elution buffer.

Half of the eluted MPRA oligo:GFP library was transformed into 200 µl NEB 10-Beta high efficiency competent cells. Transformed bacteria were split into 4 tubes, recovered in 1 ml of NEB 10-Beta/Stable Outgrowth Medium, incubated at 37 C for 1 hour. To count the CFU for a library, 10 µl of recovered cells were diluted in 10-fold serial dilution and colonies counted. The mixture was diluted to >1X10^6^ CFU/ml. To test MPRA library complexity, 12 colonies were randomly picked and inserts sequenced. To prepare the library for transfection, four 100 ml flasks of LB-Ampicillin were incubated at 37C for 6 hours. We repeated this same transformation protocol two more times, each time calculating transformation efficiency. All plasmid preparations were pooled and normalized to 1 µg/µl concentration.

### Transfection of MPRA oligo:GFP library assembly

HEK 293 cells were seeded 1.5X10^6^ cells on 2, 100 mm plate in D10 medium (DMEM supplemented with 10 %FBS) 24 hrs. before transfection. On the day of transfection, HEK 293 cells grown to 60-70% cell density were transfected with 10 µg of MPRA oligo:GFP library assembly and 20 µl of Lipofectamine 3000 (Life Technologies). The transfected cells were incubated for 24 hours in D10 medium, before rinsing with PBS. The transfection was repeated independently 5 times.

### Total RNA and DNA extraction

Total RNA and DNA were extracted using TRIzol reagent (Invitrogen). 1ml of TRIzol was added directly to the culture dish to lyse the cells and the lysate was triturated several times to homogenize. The solution was transferred to a microcentrifuge tube and incubated for 5 min. 0.2 ml of Chloroform added and mixed several times followed by incubation for 2-3 min. Lysates were centrifuged for 15 min at 12,000xg at 4C to separate the 3 phases and the aqueous phase was transferred for RNA extraction. The remaining two phases were used for DNA isolation as per the manufacturer’s protocol.

Total RNA was extracted and purified using Zymo-Spin-IC column (RNA clean& concentrator-5) including the on-column DNase digestion. A second DNase treatment was performed on the purified RNA using 5 µl of Turbo DNase (Life Technologies) in 300 µl of total volume for 1 hour at 37C followed to purify by Zymo-Spin IC column.

### cDNA, product amplification, and sequencing

First-strand cDNA synthesis was from the second DNase treated RNA using a primer specific to the 3’ GFP (RT-GFP_ 2 specific primer; Supp. Table 1) with SuperScript IV Reverse Transcriptase (Invitrogen) following the manufacturer’s protocol. Single-stranded cDNA was purified by NucleoSpin Gel & PCR clean-up column and eluted in 30 ul.

To add Illumina sequencing adaptors, we performed 8 independent 50 µl PCR reactions: 10 ng of template DNA (5 total DNA samples and 5 single-stranded cDNA), 25 µl of 2X NEBNext UltraII Q5 MM (NEB), 0.5 µM of Illumina adaptor_ GFP_R and Illumina adaptor_ GFP_F primers (Supp. Table 1) with the following conditions: 98 C for 30 sec, 10 cycles of (98 C for 10 sec, 65 C for 15sec, 72 C for 30 sec), 72 C for 1min. The PCR-amplified samples were gel-extracted to remove primers and dNTPS using NucleoSpin Gel & PCR clean-up column. Samples were sequenced by 250 bp paired ends sequencing without fragmenting the amplicons on an Illumina MiSeq (Azenta, NJ).

### MPRA data processing: Mapping of DNA and RNA barcodes

Forward and reverse fastq reads were mapped to barcodes separately using locate module of SeqKit (v2.8.0). According to the experimental design, reverse transcribed barcodes were used as a reference to align with forward reads (R1) and while forward barcodes were used a reference to align with reverse reads (R2). To reduce false positives, firstly, no mismatches were allowed for mapping. Secondly, mapped reads of length 181 were selected where barcodes were mapped to specific positions in both forward and reverse reads. Read counts for barcodes were generated separately for forward and reverse reads per replicate using custom R script (https://github.com/ankita86/MPRA_Childs_Lab). We kept the highest barcode count either from reverse or forward read for downstream analysis. We summed barcode counts per allele or element to generate aggregate counts of DNA and RNA.

### Identification of allele-specific enhancers

To identify allele-specific enhancers, paired aggregated counts of major and minor alleles or elements for both DNA and RNA was provided as an input to mpralm function of mpra bioconductor R package. This linear based model assesses the differential activity of alleles. To construct the linear model, we passed following parameters to the mpralm function: mpralm(object = mpraset, design = design, aggregate = “none”, normalize = TRUE, block = block_vector, model_type = “corr_groups”, plot = TRUE). A SNP with an FDR-adjusted p-values P<0.05 (calculated using a multiple-testing correction method to reduce the false discovery rate) is defined as an active allele-specific enhancer and those with P-value between 0.05 and 0.01 are defined as suggestive allele-specific enhancers ^21^.

### Cell lines

HEK293 (ATCC CRL-1573) and IMR-32 (gift from Dr. A. Narendran) were grown in D10 culture media (DMEM, Gibco 11995073) supplemented with 10% fetal bovine serum (FBS, Gibco 10082147) at 37°C with 5 % CO_2_. Genomic DNA was isolated using Purelink Genomic DNA mini kit (Invitrogen K182001). The STR genotype of the cells was confirmed by profiling at The Centre for Applied Genomics (TCAG) (Toronto, Canada).

### Testing rs13176921 minor allele activity using luciferase assays

Two 300 bp oligonucleotides containing either the major (A) and minor (G) alleles of rs13176921 were synthesized (Twist Biosciences; Supp. Table 1) and cloned into the pGL4.23GW-Luciferase reporter vector (Addgene). HEK 293 cells (2X10^4^ cells/100µl D10 culture medium) were seeded on white clear 96 well plate (Greiner Bio-One). The following day, 100 ng of enhancer-Luc plasmid, 5 ng of pTK-renilla as an internal control and 0.3 µl of Lipofectamine 3000 Transfection Reagent were added in 10 µl Opti-MEM (ThermoFisher) medium per well and incubated for 2 days before the medium was aspirated. 30 µl DPBS (Gibco) and 30 µl of Dual-Glo reagent (Promega) was added into each well and incubated for 30 min at room temperature. Luciferase activity was measured using Dual-Glo Luciferase SpectraMaxL (Molecular Devices) before 30 µl of Stop-Glo solution added and incubated for 30 min at room temperature and Renilla activity measured. Relative luciferase activity is luciferase activity/ Renilla activity.

### Testing NKX3.1 binding to the region around rs13176921

H1 cells expressing NKX3.1 (following ^22^) that had been differentiated to induced mural progenitor cells (iMPCs) were pelleted (kind gift from Dr. Manna Abubaker) at 760 × *g* for 5 min at 4°C, resuspended with media, cross-linked with formaldehyde to a final concentration of 1% (ThermoFisher) with shaking for 10 min at room temperature before treatment with 125 mM of Glycine (Sigma) for 5 min at room temperature. Cells were pelleted at 760 g for 5 min at 4 °C to pellet cells. Cross-linked cells were lysed with ChIP Lysis buffer (50 mM TrisHCl pH 8, 10 mM EDTA pH 8.0, 1% SDS, 1X Complete Protease inhibitor cocktail tablets (Roche), 1 mM PMSF (Sigma) for 10 min on ice. The lysate was sonicated for 25-30 cycles of 30 seconds ON, 30 seconds OFF at High Power using a Bioruptor (Diagenode). The sonicated lysate was centrifuged at 16,000 × g at 4 °C for 20 minutes and the sheared chromatin was transferred to new tubes.

Each 500 ug of sheared chromatin-protein complex was incubated with antibody targeting anti-NKX3.1 (Cell Signaling) and normal rabbit Ig G (EMD Millipore) as a negative control, respectively at 4 °C overnight. After immunoprecipitation, 30 µl of pre-adsorbed Dynabeads protein G (Invitrogen) were added and rotated further for 6 hours at 4 °C. The Antibody/protein/chromatin/bead complex was washed 4 times with RIPA buffer for 10 min at 4 °C and 2 times with TE buffer before reverse-crosslinking at 65°C overnight, while rotating using a Thermomixer at 500rpm. DNA was purified by DNA purification columns (Omega). The relative enrichment of specific regions in precipitated DNA was measured by quantitative PCR (qRT-PCR) using PowerUp SYBR green master mix (Applied Biosystems) and a QuantStudio 6 PCR machine.

### Knock out CRISPR sgRNA mutagenesis, clonal selection, and qPCR validation

All knock out CRISPR target oligos (Supp. Table 1) were designed with the ends: 5’-caccg (BbsI) N20-3’ and 5’-aaac (BbsI) N20-c-3’. Annealed oligos were inserted into pSpCas9 (BB2)-2A-GFP (pX458) vector (Addgene) ^23^digested with FastDigest BpiI (BbsI) (ThermoFisher) as above. CRISPR sgRNA plasmids were verified by sequencing. Transfection and clonal analysis was as reported ^24^.

To test knock out VCAN clones, total RNA was isolated using TRIzol reagent (Invitrogen) and purified with Zymo RNA clean and concentrator-5 (Zymo Research). Briefly, pelleted cells were lysed with 500 µl of TRIzol and incubated 5 min at room temperature. 100 µl of chloroform was added, mixed and incubated 5 min at room temperature. The mixtures were centrifuged at 12,000 g for 15 min at 4 °C, then 300 µl of aqueous phase carefully transferred to a new tube and equal volume of 100% ethanol added. For RNA purification, column purification was used (Zymo RNA clean and concentrator-5). RNA concentration was measured using a Nanodrop (ThermoFisher). For cDNA synthesis, 1 ug total RNA mixed with 4 µl of LunaScript RT mix (NEB) in a 20 µl reaction (25°C for 2 min, 55°C for 10 min at, 95°C for 1 min).

cDNA was used for real time quantitative PCR (qPCR) to determine gene expression using custom primers (Supp. Table 1). qPCR was performed using PowerUp SYBR Green Master Mix (Applied Biosystems) on a BioRad Opus 96 (BioRad). Cycle conditions followed 95°C for 3 min, 40 cycles of 95 °C for 10 s and 60 °C for 30 s. Fold increase or decrease in gene expression were determined relative to control cells using the ΔΔCT method.

### Statistical analysis

All experimental replicates are noted in the results, and are typically a minimum of 3 biological replicates, and averaging 3 technical replicates (qPCR). Experimental statistics were calculated in GraphPad Prism using tests noted in figure legends.

### Data and code availability

The datasets generated during this study are available in the Supplemental Tables. The sequencing data is deposited in GEO with the accession: GSE316826. The code generated during this study is available at https://github.com/ankita86/MPRA_Childs_Lab.

## Results

### A multiplexed reporter assay identifies enhancers

In vitro Multiplexed Reporter Assays (MPRAs) are used for unbiased detection of allele-specific enhancer activity. We modified the original MPRA design^20^ to analyze uncharacterized stroke-associated variants from GWAS studies. For this medium-scale screen, we selected 44 variants in 8 human genes associated with stroke (and subtypes), SVD MRI-markers, vascular risk factors (HDL- and LDL-cholesterol, triglycerides) or related diseases, such as venous thromboembolism or coronary artery disease (Supp. Table 3)^3,9,11,12,24,25^ The variants are located in intergenic or intronic regions adjacent to the following genes: *EGF Containing Fibulin Extracellular Matrix Protein 1/Fibulin 3 (EFEMP1), Forkhead Box F2 (FOXF2), 3-Hydroxyanthranilate 3,4-Dioxygenase (HAAO), Potassium Two Pore Domain Channel Subfamily K Member 3 (KCNK3), N-Myristoyltransferase 1 (NMT1), OPA1 Mitochondrial Dynamin Like GTPase (OPA1), RAS Like Family 12 (RASL12), Signal Transducer And Activator Of Transcription 3 (STAT3)* and *Versican (VCAN)*.

A pool of 968 oligonucleotides (Supp. Table 4), each 293 bp in length, were synthesized. The oligo design contains forward adaptors, 204 bp of genomic context with major or minor alleles at the center, and barcodes (Fig. 1). Each oligo was synthesized with 1 of 11 different barcodes to avoid any sequence bias in barcoding. The oligo library was cloned upstream of a minimal promoter and GFP (Fig. 1). In this way, any enhancement or repression of GFP expression levels can be quantitated through comparison of barcodes from DNA (transfection efficiency) and RNA for the same barcode (expression levels). Correlated expression 11 oligos with the same core sequence but different barcodes provide built-in redundancy (Supp. Table 4). As a positive control, we included known allele-specific enhancer variants in *FOXF2* we have previously validated using luciferase, ChIP and CRISPR assays^24^.

**Figure 1:**
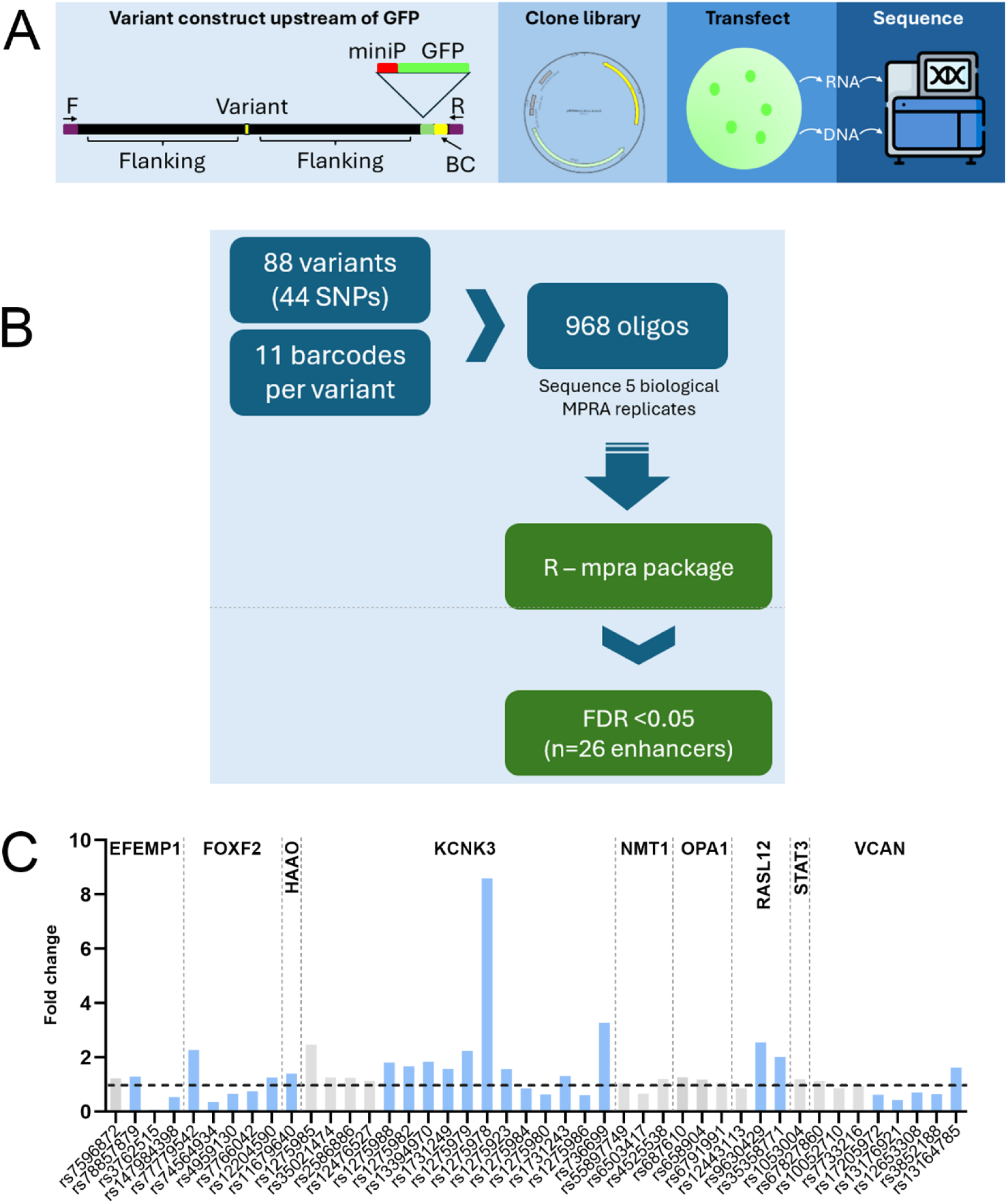
MPRA methodology. A) Schematic of the MPRA synthetic oligonucleotide including a forward primer (F), 102 bp of flanking sequence before, and 102 bp of flanking sequence after, the variant followed by a miniPromotor (miniP)-GFP construct, a 10 bp barcode and reverse primer (R). The oligo library was cloned into the pMPRAv3:Δluc:ΔxbaI vector and transfected into HEK293 cells. DNA and RNA was isolated separately and sequenced using MiSeq. B) Bioinformatic workflow for the MPRA analysis showing how allele-specific enhancers were identified. C) Fold change in expression for alelle specific enhancers in the indicated genes. Blue bars show significant allele-specific activity.

As proof of concept that the experimental design could detect enhancer activity, GFP expression in transfected cells confirmed the varied ability of the oligos to confer enhancer activity (Supp. Fig. 1). DNA and RNA were isolated from 5 biologically independent replicate transfections (Fig. 1A). To ensure that sequencing reads were from transfected material and not endogenous genomic DNA and RNA, we sequenced an amplified fragment containing the 3’ end of GFP and the barcode bracketing the test oligo sequence. The sequence was bioinformatically trimmed, and non-specific amplification products that were not exactly 181 bp were removed. After this quality control step, an average of 83.62% of the unique reads were mapped in both DNA and RNA samples (Supp. Table 5). The raw counts from the five RNA and DNA samples show correlation of between 0.99 and 1.00 at the DNA level and 0.97 and 1.00 at the RNA level, suggesting high concordance between transfections (Supp. Fig. 2, Supp. Fig. 3). The majority of the 968 barcodes were represented in the sequencing. Only 21 (2.17%) were completely lacking from DNA and RNA, while 146 were found at low count levels (less than 5 reads in both DNA and RNA). This suggests that the diversity of the library was well represented in the transfections (Supp. Fig. 4).

We next identified allele specific enhancers using Bioconductor mpralm ^26^. We discovered 26 allele specific enhancers that met a significant false discovery rate (FDR) of <0.05, and a further 4 allele-specific enhancers that meet a suggestive FDR of P<0.1 (Supp. Table 6). In summary, MPRA analysis of stroke- and SVD-associated variants identified novel allele-specific enhancers suggesting potential functions in transcriptional regulation of the underlying genes.

### Previously identified FOXF2 variants are allele specific enhancers in the MPRA

To confirm that the MPRA identifies bone fide enhancers, we included previously characterized variants in *FOXF2* that are associated with CSVD. We previously showed that a switch from C to T in rs74564934 results in a ~50% reduction in enhancer activity using a cell-based assay^24^. We also included rs12204590, a lead variant associated with all stroke ^16^ but where we did not identify allele-specific effects in previous cell-based assays. The MPRA shows that rs74564934, rs4959130 and rs7766042 show significant allele-specific activity, with the minor allele decreasing expression, agreeing with our experimental data (Fig. 2). In agreement with our published work, rs12204590 does not show allele-specific activity. To extend this work, we predict that multiple transcription factor sites are predicted to be lost and gained at each of the SNVs using the MotifBreakR algorithm which looks for predicted transcription factor binding sites that are disrupted by the SNV ^27^ (Supp. Fig. 5). Thus, the MPRA using *FOXF2* variants is consistent with experimentally validated data, suggesting it can be applied to identify activity of unknown variants.

**Figure 2:**
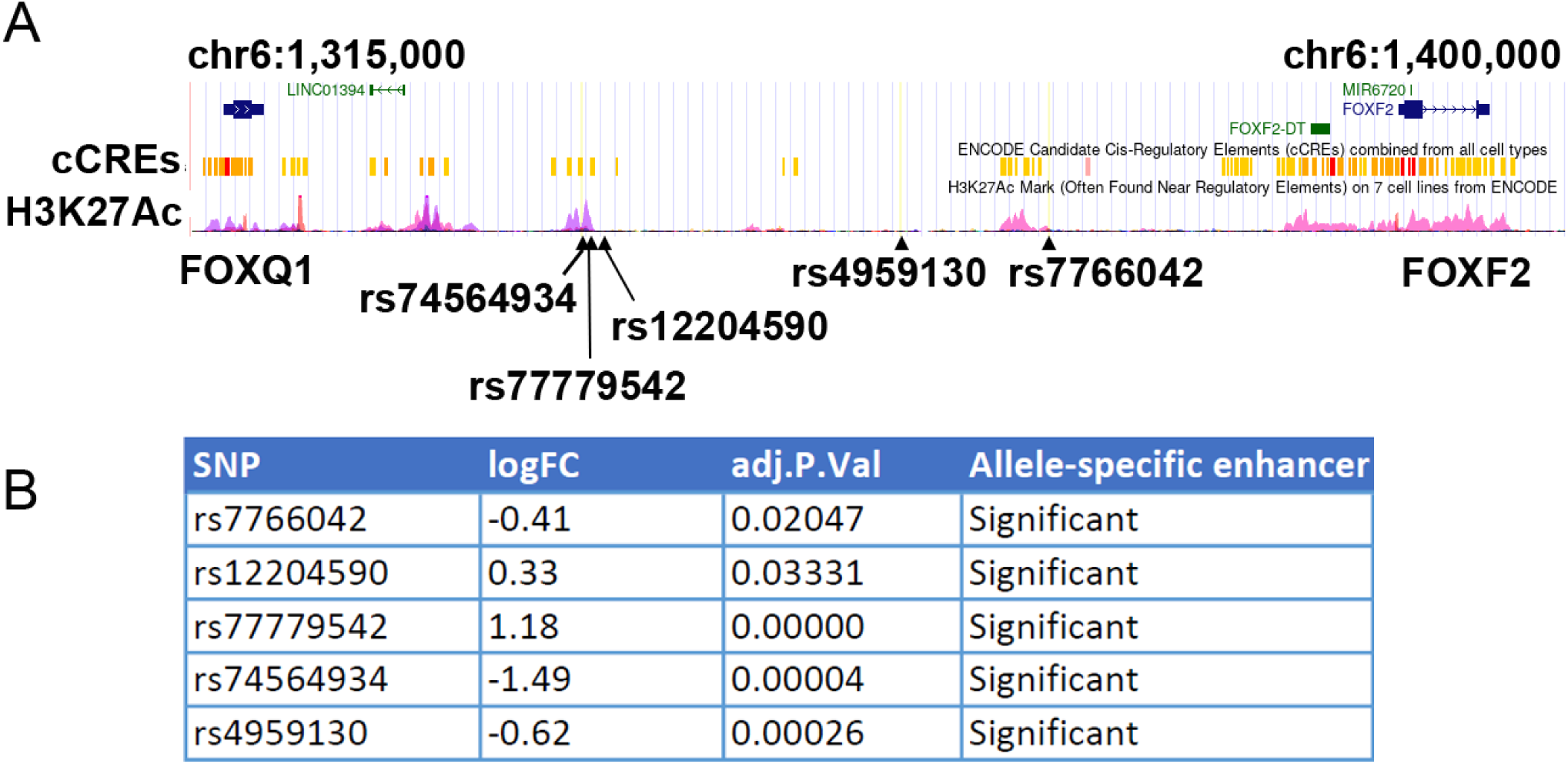
The MPRA identifies known allele-specific enhancers in FOXF2. A) Intergenic region between FOXQ1 and FOXF2 showing location of variants that are significant allele-specific enhancers from the MPRA. B) Table showing the log fold change and adjusted p value for the expression difference between major and minor alleles (allele-specific enhancers).

### Novel allele-specific enhancers in other CSVD and Stroke-associated genes

We tested for allele-specific for several additional stroke-associated genes. We find that *EGF-Containing Fibulin-Like Extracellular Matrix Protein 1 (EFEMP1*, also known as *Fibulin-3*) has 3 allele-specific enhancers in the MPRA (Supp. Fig. 6). Rs376525 strongly decreases expression in the MPRA, and several transcription factor sites are predicted to be lost and gained at each of the SNVs using the MotifBreakR algorithm (Supp. Fig. 7).

One SNV near the 3-hydroxyanthranilate 3,4-dioxygenase (HAAO) gene was tested in our assay and had significant allele specific enhancer activity (Supp. Fig. 8). HAAO is modestly lowly expressed in developing mural cells, but higher in adult mural cells macrophages of the brain, and the kidney and arteries of adults. The rs11679640 SNV is predicted to strongly decrease binding of multiple retinoic acid orphan receptors, that act as transcription factors (ROR-α, -β, and -γ, encoded by *RORA*, *RORB*, and *RORC* respectively (Supp. Fig. 9)).

In the MPRA, we identified 12 allele-specific enhancers for *KCNK3* that either increase or decrease its expression (Supp. Fig. 10). 10 of the 12 allele-specific enhancers increase reporter expression in our assay suggesting that they may be located in transcriptional repressors, especially rs1275978 and rs1275979 that have strong expression differences. Several transcription factor sites are predicted to be lost and gained at each of the SNVs using the MotifBreakR algorithm (Supp. Fig. 11).

*RASL12* is poorly studied, but is expressed in vascular mural cells, arteries, and vascular smooth muscle cells in human sequencing databases (Supp. Fig 12). *RAS Like Family 12 (RASL12)* has two allele-specific enhancers through our analysis that increase reporter expression (Supp. Fig. 12). Although rs12449945, an intronic variant within *RASL12,* is associated with white matter hyperintensity burden volume^11^, the presence of the minor allele showed no significance in our study. Several transcription factor sites are predicted to be lost and gained at each of the SNVs using the MotifBreakR algorithm (Supp. Fig. 13).

Three other genes tested, *NMT1, OPA1 and STAT3,* had no allele-specific enhancers in our MPRA.

### Novel allele-specific enhancers in VCAN

Versican is a chondroitin sulfate proteoglycan (CSPG2) with roles in primordial ECM deposition in development. In zebrafish single cell sequencing, *vcana* and *vcanb* (two orthologs of human *VCAN*) are expressed in developing vascular mural cells (Fig. 4). In the first trimester developing human brain, *VCAN* is expressed in vascular mural cells. In adult brains, *VCAN* is expressed in both vascular cells, arteries, and brain oligodendrocyte precursor cells, as well as arteries (Fig. 4). Thus, *VCAN* is expressed in locations consistent with a role in CSVD.

We tested 8 *VCAN* variants in LD (r^2^ >0.88) with the lead variant rs67827860 associated with WMH and NODDI in GWAS studies in *Versican (VCAN)*. All variants are located in intron 12 of *VCAN*. The MPRA shows that four variants have significant allele-specific enhancer activity (rs17205972, rs13176921, rs12653308, and rs3852188), while rs13164785 is suggestive (Supp. Table 4; Fig. 3). Our analysis therefore identifies novel intronic enhancers for *VCAN*. Several transcription factor sites are predicted to be lost and gained at each of the SNVs using the MotifBreakR algorithm, including the transcription factor NKX3.1 (Supp. Fig. 14).

**Figure 3:**
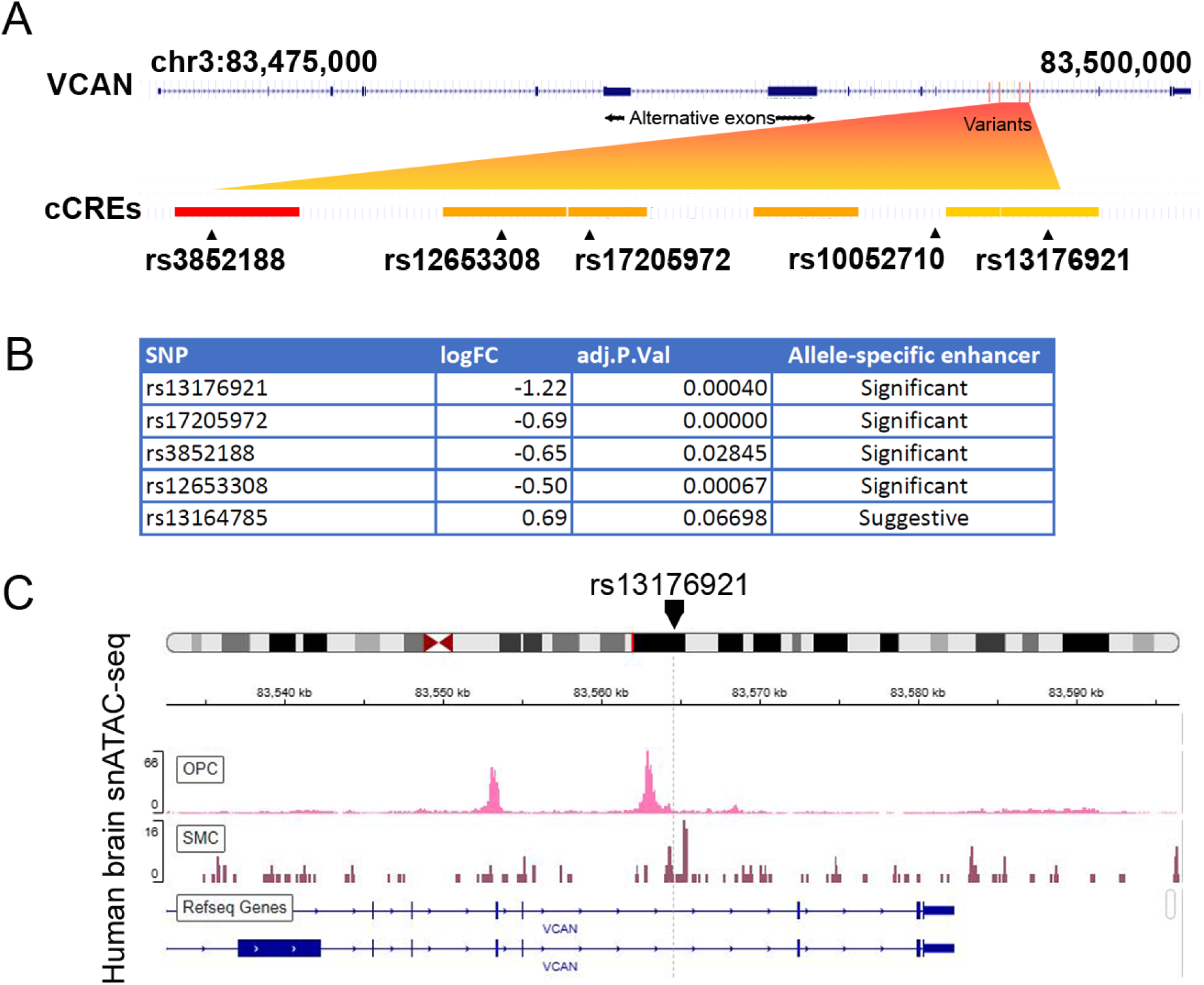
The MPRA identifies allele-specific enhancers in VCAN. A) Map of the VCAN gene with enlargement of the region of the variants identified in our study. The orange trapezoid shows how variants map to intron 12 of VCAN, in Candidate Cis-Regulatory Elements (cCREs) from ENCODE (red and yellow bars). B) Table showing log fold change an adjusted P value as determined by the MRPA assay for significant allele-specific enhancers. C) Representative human brain snATAC-seq data from CATlas^1^ around rs13176921 in oligodendrocyte precursor cells (OPC) and smooth muscle cells (SMCs) showing open chromatin in the region of this variant.

**Figure 4:**
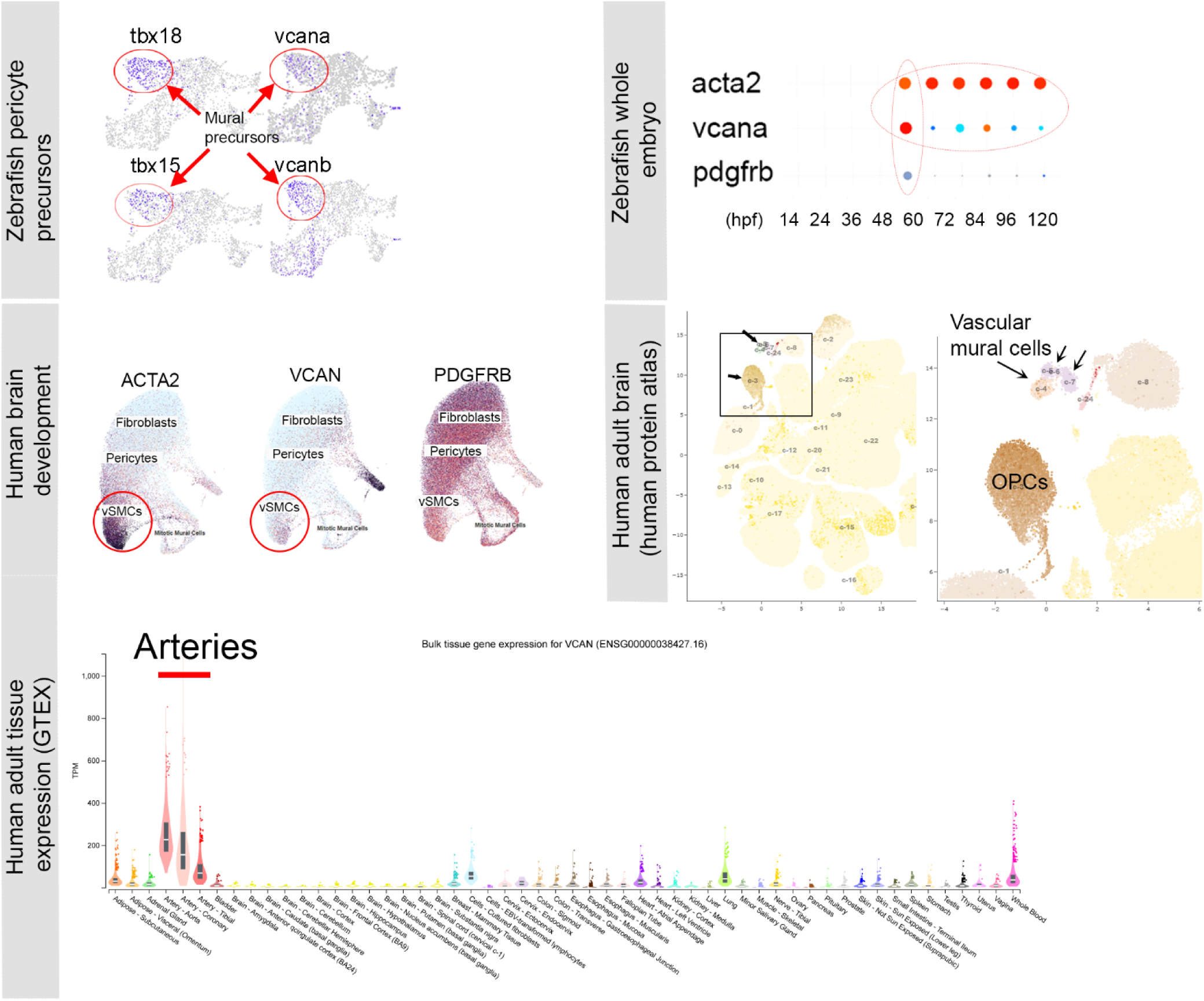
Expression of Versican mRNA in zebrafish and human. In zebrafish, *versican-a (vcana)* and *versican-b (vcanb)* are both expressed in FACs-sorted pericyte precursors^2^. In zebrafish whole embryo, *vcana* is co-expressed in *acta2*-expressing vSMCs and 60 hpf *pdgfrb*-expressing pericytes^3^. In human first trimester sorted vascular cells from the brain, *VCAN* is co-expressed with both *ACTA2* and *PDGFRB*^4^. In the human adult brain, VCAN is strongly expressed in oligodendrocyte precursors (OPCs), and in vascular mural cells, especially fibroblasts^5^. In human adult tissues from the GTEX database, VCAN is strongly expressed in the aorta and arteries^6^.

### Allele-specific enhancers in VCAN do not alter alternative splicing

*VCAN* is a highly spliced transcript, with major splice forms V0, V1, V2, and V3. It is possible that disease-associated variants could affect splicing, even though all genomic variants tested in our MPRA are in an intron downstream of the alternative exons, and it is unlikely. We used qPCR to assay *VCAN* splice form expression in IMR-32, a neuroblastoma line robustly expresses all V0-V3 variants. We note that the SNVs in IMR-32 are of the major allele in this line, and this is the only cell line we identified with expression of all *VCAN* splice forms (Supp. Fig. 15.). To test intron 12 variant effects on splicing vs. gene expression, we deleted a ~2 kb intronic region containing 3 significant allele-specific enhancers in IMR-32 cells by simultaneous transfection of two CRISPR guides and Cas9 (Fig. 5A). We single-cell cloned two CRISPR modified lines and sequenced breakpoints around the deletion, confirming they both had similar sequences (Fig 5B). Using qPCR, we show that *VCAN* V3 expression is significantly reduced when the 2kb enhancer is deleted (Fig 5C), suggesting that the 2 kb region normally enhances *VCAN* expression. However, there were no changes to alternative splicing of *VCAN* V0, V1 and V2 using qPCR. Because there is significant downregulation of V3 (total *VCAN* expression) in the enhancer-deleted lines, our data suggests that intron 12 promotes *VCAN* expression but does not modulate splicing (Fig 5C).

**Figure 5:**
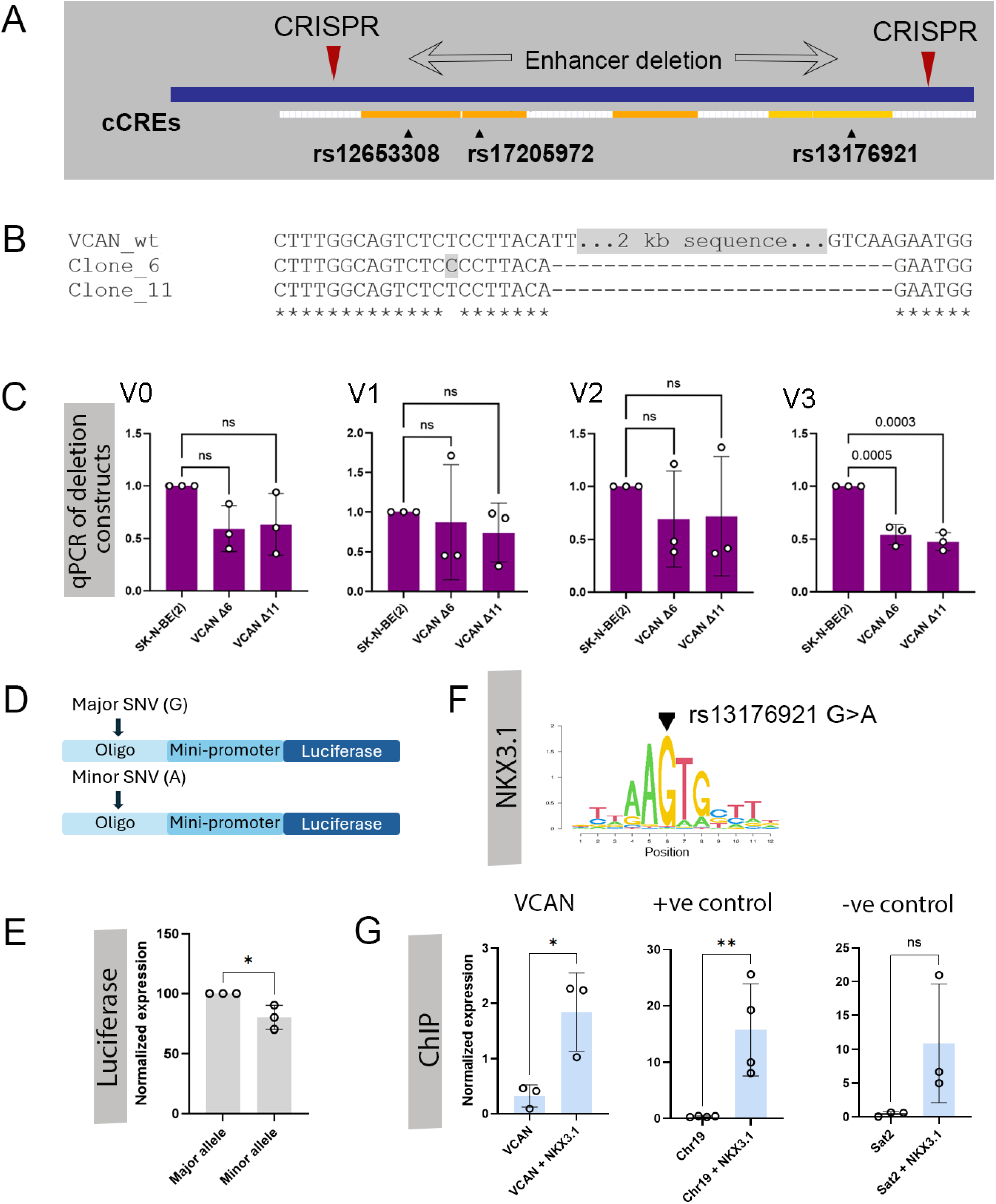
Modulation of a novel VCAN enhancer leads to decreased expression. NKX3.1 binds to the region around rs13176921. A) Schematic of where a 2 kb intronic enhancer region was deleted using two CRISPRs. B) Sequences of independent clones with the 2 kb region deleted. C) qPCR of endogenous VCAN in the deleted clones showing that VCAN V3 is significantly downregulated when the enhancer is deleted but there are no significant changes in V0, V1 and V2. D) Schematic of constructs with the major and minor alleles of rs13176921 in a 300bp oligo. E) Luciferase activity is decreased when the minor allele is present, relative to the major allele. N=3 biological replicates, with 3 technical replicates. F) Position weight matrix for NKX3.1 from JASPAR showing location of rs13176921 SNV. G) ChIP for NKX3.1 in the region of rs13176921, a positive control genomic region known to bind NKX3.1 (Chr 19) and a negative control region (Sat2). Statistics used a One Way ANOVA with Dunnett’s multiple comparison test, or the Student’s t-test.

### Identification of a functional SNV in intron 12 of VCAN

In the 2kb intron 12 region we deleted there are 3 disease-associated SNVs. rs10052710 is associated with the Neurite Density Index (NDI) metric ^13^, rs17205972 is associated with WMH volume ^11^, and rs13176921 is associated with NDI in four different brain regions ^13^. However, allele-specific enhancer activity was non-significant for rs10052710, and while it was significant for both rs17205972 and rs13176921. rs13176921 shows a larger magnitude of fold change between major and minor alleles. Using SNP2TFBS ^28^ and MotifBreakR ^27^ to predict transcription factor binding sites disrupted by rs13176921, a strong binding site for the transcription factor NKX3.1 was identified where the G>A change is at a highly conserved position, and predicted to abolish binding in the minor allele (Fig 5F). NKX3.1 is a transcription factor that powerfully induces a mural cell precursor phenotype, which intrigues us because of the relationship between mural cells, ECM and vascular stability ^22,29^. rs13176921 is located in a region of open chromatin as shown by snATACseq in the CATlas ^30^ (Fig 3C), in both oligodendrocyte precursor cells and vascular smooth muscle cells. To specifically test that the SNV (G-A) changes enhancer expression we used a luciferase assay, cloning two identical 300 bp oligonucleotides, varying only at the SNV position, ahead of a minimal enhancer and luciferase (Fig 5D). This showed that the 300 bp region around rs13176921 is a very strong transcriptional enhancer. Further, presence of the minor allele (G), as opposed to the major allele (A), significantly reduces luciferase activity confirming the results of the MPRA assay in an independent assay (Fig 5E). Finally, show that NKX3.1 binds to this region using Chromatin Immunoprecipitation (Fig 5G), suggesting it could regulate this enhancer sequence.

## Discussion

Disease-associated SNVs are a window into the function of the genome. GWAS coupled with bioinformatic analysis have advanced substantially but provide incomplete insight into variant function. In this study we use a medium-throughput assay to functionally test GWAS SNVs in genes associated with stroke and CSVD. Of the 44 variants tested in our MPRA assay, 26 had allele-specific enhancer properties suggesting that disease-associated variants arising out of these highly powered GWAS studies identify DNA sequences capable of modulating gene expression, sensitive to a single SNV.

We included positive control variants from our previous analysis of *FOXF2* stroke associated variants^24^. These variants had been tested using cell-based assays that showed that constructs harboring the minor allele of rs74564934 had ~50% expression of constructs with the major allele. We previously identified that there was decreased binding of the transcription factor ETS1 to this variant as the variant is located within the ETS1 site ^24^. We extend this work here by identifying additional enhancers in the region around adjacent variants rs7766042 and rs4959130. Both variants are allele-specific enhancers in our current MPRA assay. The three *FOXF2* variants are spaced 10-20 kb apart and each is in a different ENCODE CCRE, suggesting complex regulation of *FOXF2* from multiple enhancers in this intergenic region. Thus, we both confirmed and extended previous results using the MPRA.

We focus on *Versican* for follow-up due to the intriguing association of some variants with SVD MRI-markers in the elderly but also in young adults ^11,13^. We provide multiple lines of evidence that intron 12 of *VCAN* has at least 4 transcriptional enhancers that positively regulate *VCAN* expression. Presence of the minor (risk) variant for 4 SNVs decreases reporter expression in vitro. Two assays in cells (deletion of the enhancer, and luciferase reporter assays) demonstrate the functional role of the major allele of rs13176921 in positively regulating *VCAN* expression. Further, we demonstrate the transcription factor NKX3.1 binds to the enhancer surrounding Rs13176921. NKX3.1 binding is predicted to be completely abolished by presence of the minor allele. Rs13176921 is also predicted to bind to other transcription factors such as DNMT1, but binding is not predicted to be disrupted using MotifBreakR and therefore was not studied further here. However additional testing of other factors binding in the region to understand enhancer architecture would be valuable.

rs13176921 is present globally in an average of 23% of humans (dbSNP). Versican is a component of the matrisome and is an ECM protein. Developmental ECM is essential for establishing the mechanical and signaling environment for cells. Many ECM proteins have long half-lives and are largely synthesized largely in development, setting up initial patterns of vascular stability for the lifespan. Versican, together with hyaluronan, a highly hydrated polysaccharide, forms a soft ‘provisional matrix’ ^31^ in development. Versican is a large protein that can be modified by chondroitin sulfate proteoglycan on alternatively spliced exons present in the V0, V1 and V2 splice forms. The core protein splice variant, V3, is not glycated and is expressed in provisional matrix promoting tropoelastin synthesis and elastin deposition ^32,33^. Non-elastic, stiff blood vessels cannot respond to changes in pulsatile blood flow, resulting in hypertension, a risk factor that enhances SVD. *VCAN* is also expressed in vascular mural cells and oligodendrocytes in developing and adult humans and oligodendrocytes after injury ^34^. The strongest expression on GTEX is in the aorta, and arteries, consistent with a vascular role for *VCAN*.

We assayed SNVs in other stroke and CSVD-associated genes. *EFEMP1 (Fibulin 3)* is a secreted glycoprotein expressed by connective tissues and interacts with elastic fibers. While EFEMP1 binds the EGFR receptor, it also has roles in cell migration and adhesion, along with roles in the formation and homeostasis of elastic tissues. *EFEMP1* is strongly expressed in arteries and EFEMP1 binds to vascular smooth muscle cells. *EFEMP1* is processed by *HTRA1* (the CADASIL2 gene causative of SVD^35,36^). EFEMP1 is a member of the matrisome group of ECM proteins and appears to be a strong candidate for follow up for a causative role in SVD. We found 3 allele specific enhancers for *EFEMP1* in our MPRA.

HAAO is an intracellular dioxygenase involved in the synthesis of quinolinic acid. Rs11679640 was identified as a genome-wide significant locus for WMH intensity burden ^37^. Quinolinic acid, is involved in neuroinflammation, suggesting how changes in HAAO expression might lead to consequences in the brain. HAAO expression has not previously been reported in mural cells of the brain but human protein atlas data suggests it is enriched in mural cells and macrophages. MotifBreakR predicts that the minor allele of Rs11679640 will strongly reduce the binding of retinoic acid receptors; this prediction has not yet been experimentally validated.

Intergenic and intronic variants near the *Potassium Two Pore Domain Channel Subfamily K Member 3 (KCNK3)* are associated with cerebrovascular events in a multiancestry study ^38^, with all stroke^25^ and with blood pressure. Loss of *KCNK3* is associated with pulmonary arterial hypertension and increased proliferation, vasoconstriction, and inflammation in smooth muscle cells ^39^. KCNK3 is a potassium channel active in pulmonary smooth muscle cells, resulting in autosomal dominant pulmonary hypertension when mutated^40^. In human tissue single cell sequencing, *KCNK3* is most highly expressed in smooth muscle cells (Supp. Fig 8). Interestingly, 9 of the *KCNK3* variants we tested significantly increased enhancer activity, one as high as 8-fold. GTEX shows that minor alleles of many of the tested SNVs including rs1275985, rs1275986, rs1275923, rs1275978, rs1275979, and rs1275980, for example, have higher expression of transcripts that have the minor allele in multiple tissues including pancreas and heart, agreeing with our data that the minor alleles of these SNVs may increase activity of the enhancer.

RASL12 is a poorly studied small Ras-like GTPase previously expressed in mural cells and shown to block YAP1 nuclear localization^41^. *RASL12* binds *RASSF1A*, recruiting it to microtubules ^41^. As *RASSF1A* interacts with Hippo signaling, it is possible that *RASL12* may act in smooth muscle cells via Hippo pathway, coupling Ras signaling to the Hippo pathway. Hippo signaling is needed for vascular smooth muscle cell differentiation ^42^. *RASL12* has strong expression in human embryonic brain vascular mural cells, human adult vascular mural cells (and astrocytes), along with strong expression in human arteries suggesting a plausible role in SVD. We identified 2 intronic allele-specific enhancers in *RASL12*.

While GWA studies identify a large number of variants, our MPRA has identified variants with cis-regulatory enhancer activity vs. variants that are simply in LD with important variants. This information is important for understanding risk in individuals, particularly for those who would benefit from preventative therapies who have a comorbidity such as hypertension. Hypertension increases the burden of neurological damage in individuals who carry high risk SVD-associated variants^11^. Theoretically, the long latency between detectable brain changes on MRI in young adults, and SVD and stroke in later life gives a window to offer lifestyle changes and therapeutics for those at greatest risk.

### Limitations

Although we are able to identify transcriptional activity using the MPRA, a limitation of the assay design is that it uses a relatively short oligo. Typical mammalian cis-regulatory enhancers are 100-1000 bp in length ^43^ which may exceed (or be shorter than) the length of the oligonucleotide used in the MPRA. Further, enhancers bind groups of transcription factors that require ‘collective occupancy’ ^44^ which is not accounted for in the agnostic design of the MPRA that only considers the SNV placed at the center of the oligo. Enhancers may have multiple transcription factor binding sites that are interdependent in their binding, and the oligo may not include these. Enhancer activity *in vivo* may also require trans long-range interactions with promoters and transcriptional machinery elsewhere in the chromosome. The MPRA removes the enhancer from long-range interactions, is not subject to epigenetic control, and is often run in a cell line that does not exactly match the context of gene expression due to the need for high transfection efficiency of a library ^15^. Nonetheless, the speed and breadth of the MPRA can rapidly identify functional SNVs for further dissection with functional studies in an appropriate genomic context.

The large number of variants associated with stroke and CSVD from GWAS, and their lack of conservation of variants, even among mammals, makes testing variants *in vivo* difficult. The MPRA assay was done in HEK cells, due to the need for high transfectability of a large library of variants. Possible sources of error in our study include the lack of tissue-specific factors for gene expression in this cell line. However, as we detected general enhancer activity from most enhancers tested, HEK was a reasonable choice for the experiments. For in vitro validation of the *VCAN* variants, only the neuroblastoma IMR-32 cells expressed all *VCAN* splice forms at levels sufficient for testing whether variants affect its splicing. This cell line was therefore chosen for biological validation. Development of an in vivo MPRA assay would be ideal, but this is not currently available for vascular cells.

## Appendices

### List of Tables

Supp. Table 1: Primer sequences

Supp. Table 2: Chemicals and Regents

Supp. Table 3: MPRA Oligonucleotides synthesized

Supp. Table 4: Genes and associated variants tested in the MPRA assay

Supp. Table 5: Mapping of barcodes for each replicate

Supp. Table 6: Allele specific enhancers

**Supplemental Figures 1-14**.

## Acknowledgements

Salary support for bioinformatic analysis (AN) is supported by the Alberta Children’s Hospital Research Institute.

