## Supplemental figures for "Characterization of variants associated with Cerebral Small Vessel Disease identifies a functional SNV in Versican"

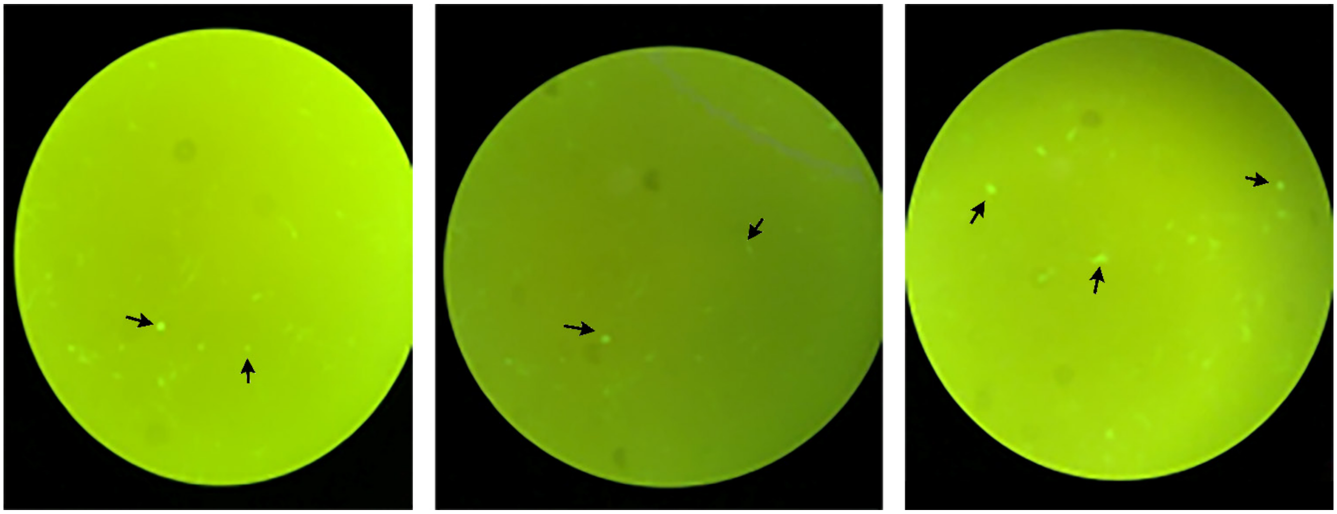

**Supplemental Fig 1: GFP expression in transfected cells.**

Example of variable GFP expression in transfected cells prior to FACS sorting. Colonies with higher GFP expression higher than baseline suggest these colonies have strong enhancer activity.

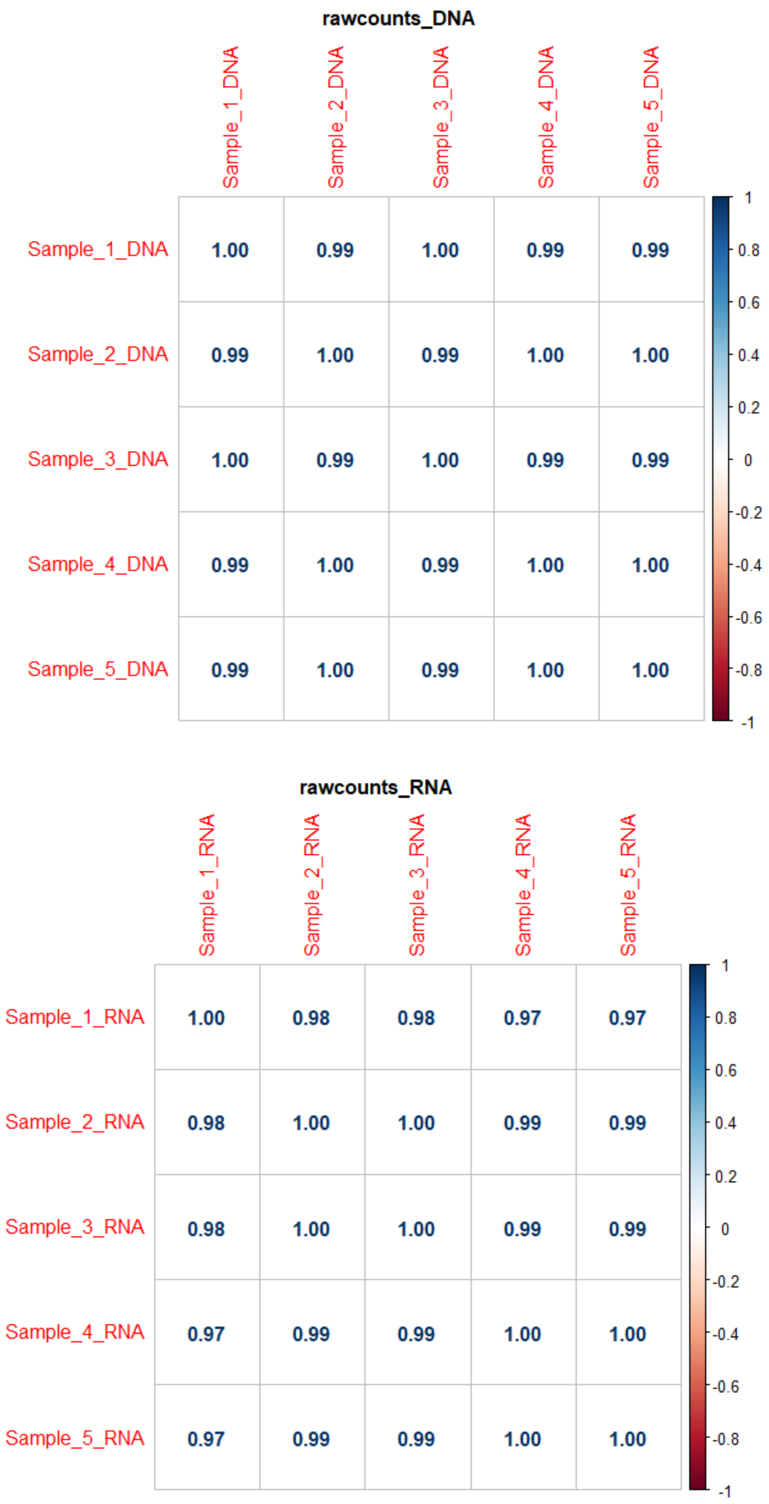


**Supplemental Fig 2: Correlation among the raw counts for DNA and RNA across replicates** suggests high correlation between aggregated counts among replicates.

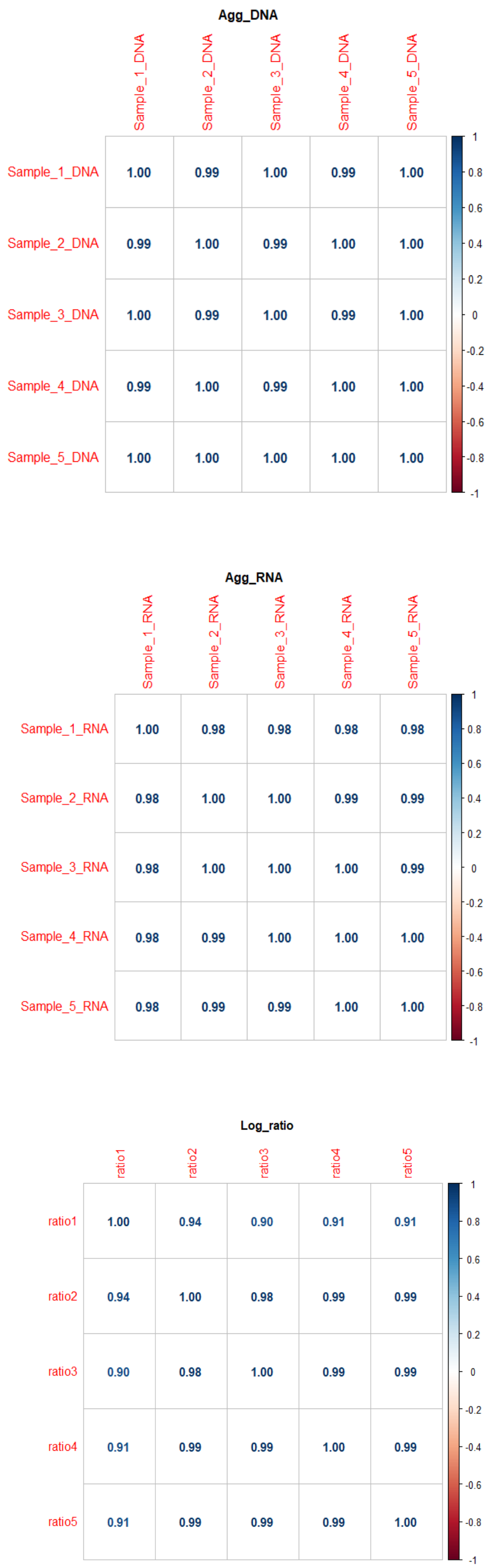


**Supplemental Fig 3: Correlation between replicates for aggregated raw counts of DNA, RNA and their Log ratios i.e. (log2(RNA/DNA)** shows high correlation between aggregated counts among replicates

| | 10 samples |
|---|---|
| Barcodes | 968 |
| Counts=0 in all replicates (both DNA and RNA) | 21 (2.17%) |
| Counts <5 in more than 5 samples (both DNA and RNA) | 146 (15%) |

**Supplemental Fig 4: Representation of barcodes in the sequenced library (5 DNA and 5 RNA replicates)**

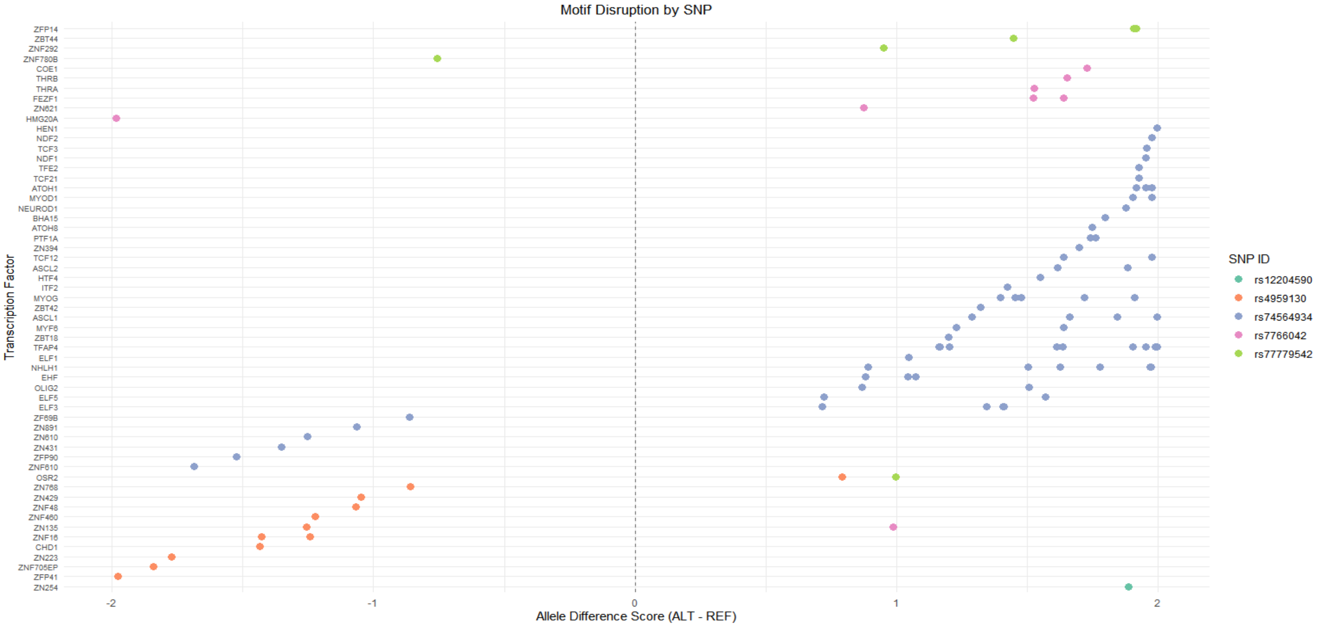


**Supplemental Fig 5: Predicted effect of SNV on transcription factor binding from MotifBreakR [7] for FOXF2 SNVs.**

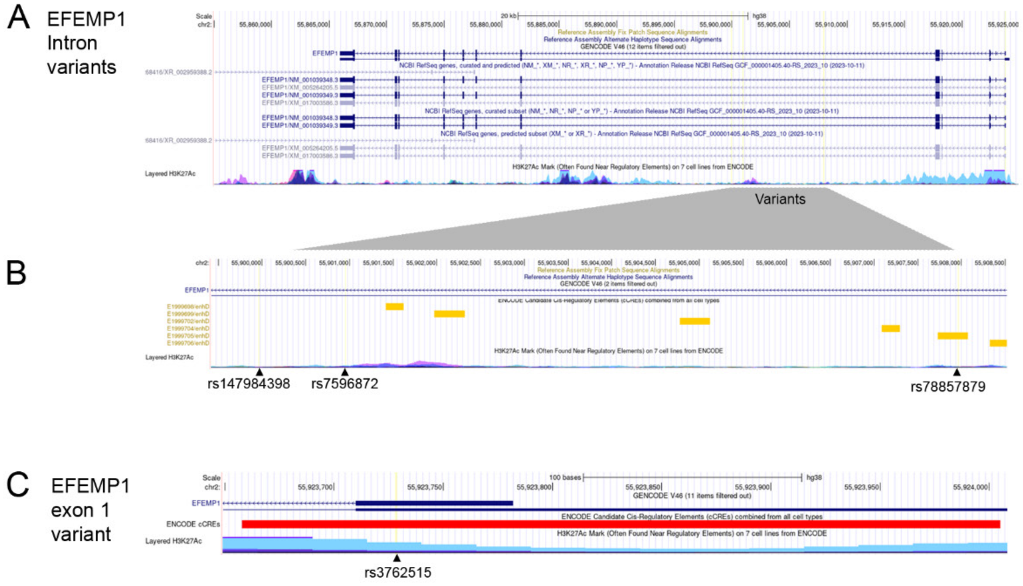


D

| SNP | logFC | adj.P.Val | Significant | Gene |
|---|---|---|---|---|
| rs7596872 | 0.3002199 | 0.487998552 | NS | EFEMP1 |
| rs78857879 | 0.3762663 | 0.000466802 | Significant | EFEMP1 |
| rs3762515 | -3.8411093 | 0.028449814 | Significant | EFEMP1 |
| rs147984398 | -0.887799 | 0.093239736 | Suggestive | EFEMP1 |

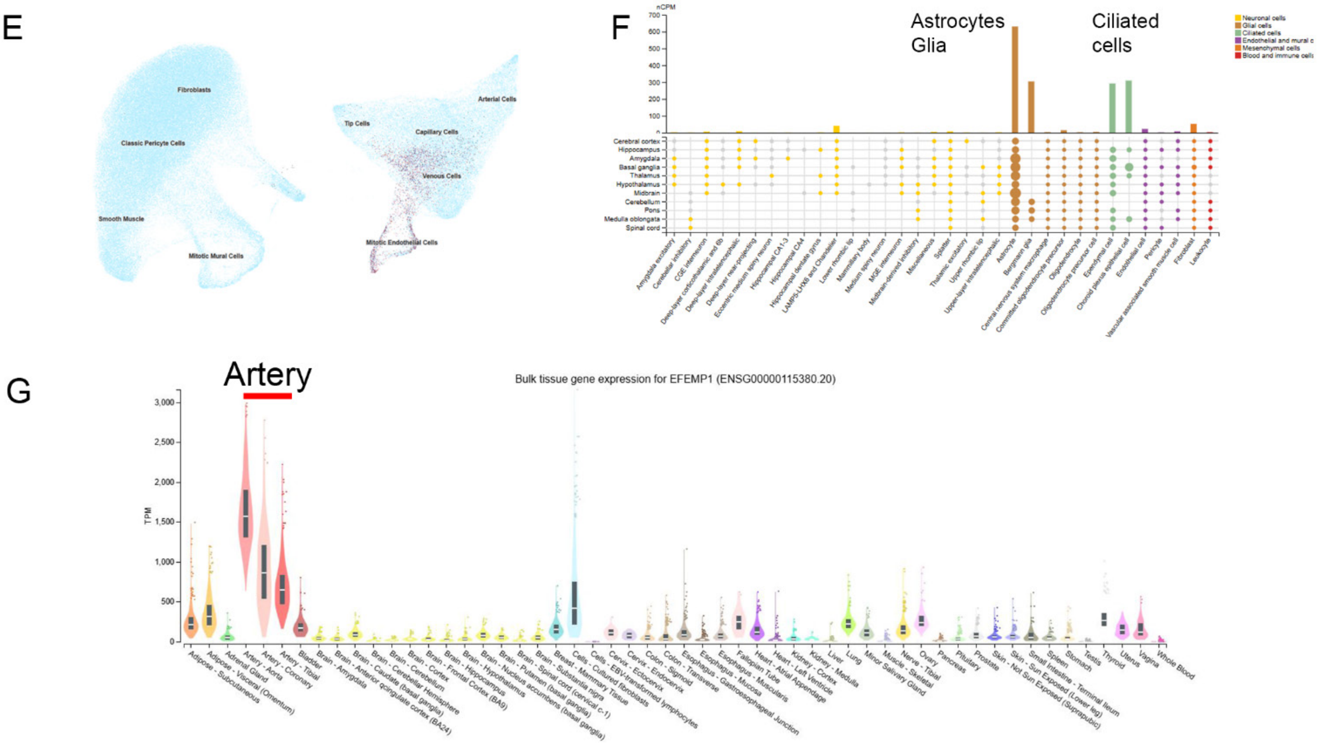


**Supplemental Fig 6: EFEMP1 variant locations**

A) Map of the EFEMP1 gene with enlargement of the region of the variants identified in our study. The grey bar shows how variants map on to intron 7 of the gene. B) Some variants are located in Candidate Cis-Regulatory Elements (cCREs) from ENCODE (red and yellow bars, the variant is indicated below. C) one variant is located in exon 1. D) Table showing the log fold change and adjusted p value for allele specific enhancers as determined by the MRPA assay. (NS means not significant). E) Low expression of EFEMP1 in human developing brain vessels[4]. F) Expression in astrocytes, glia and ciliated cells in human single nucleus adult brain sequencing[5]. F) Expression in arteries in the GTex dataset[6].

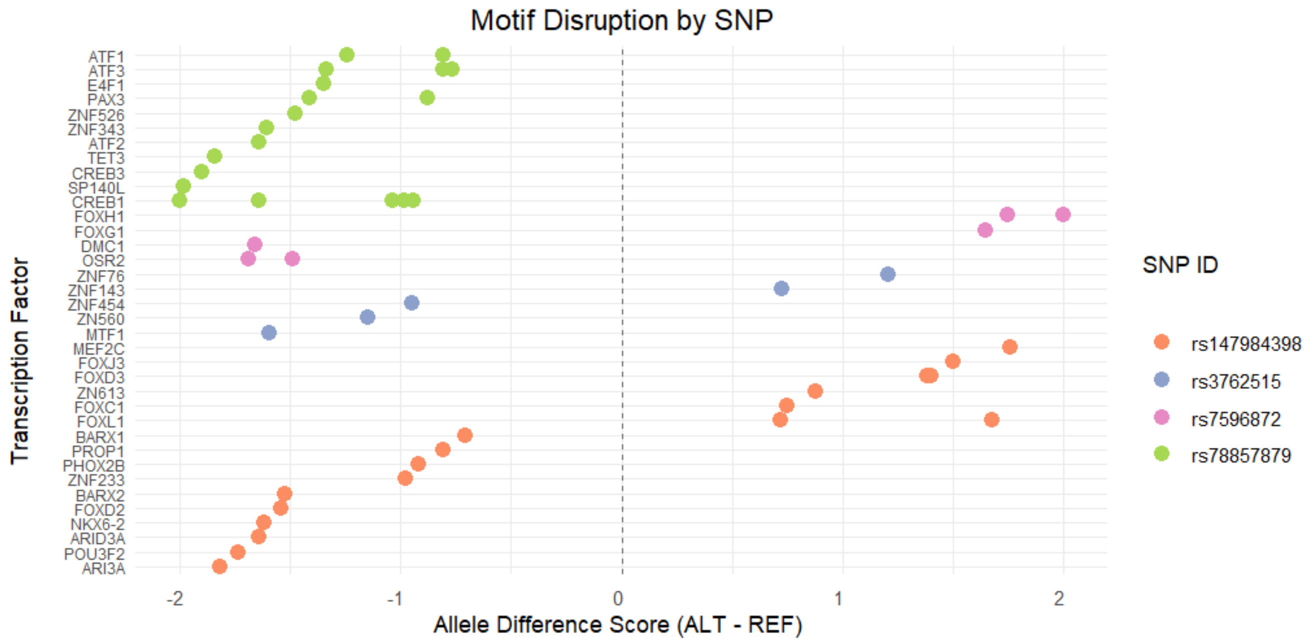


**Supplemental Fig 7: Predicted effect of SNV on transcription factor binding from MotifBreakR [7] for EFEMP1 SNVs**

A

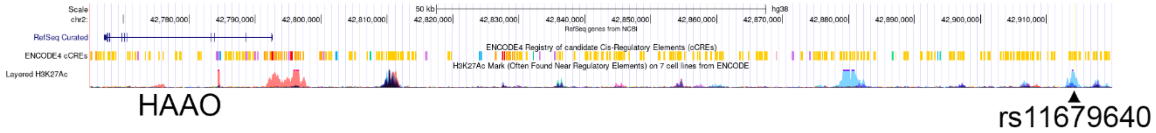


B

| SNP | logFC | adj.P.Val | Significant | Gene |
|---|---|---|---|---|
| rs11679640 | 0.481486 | 1.32134E-05 | Significant | HAAO |

C D

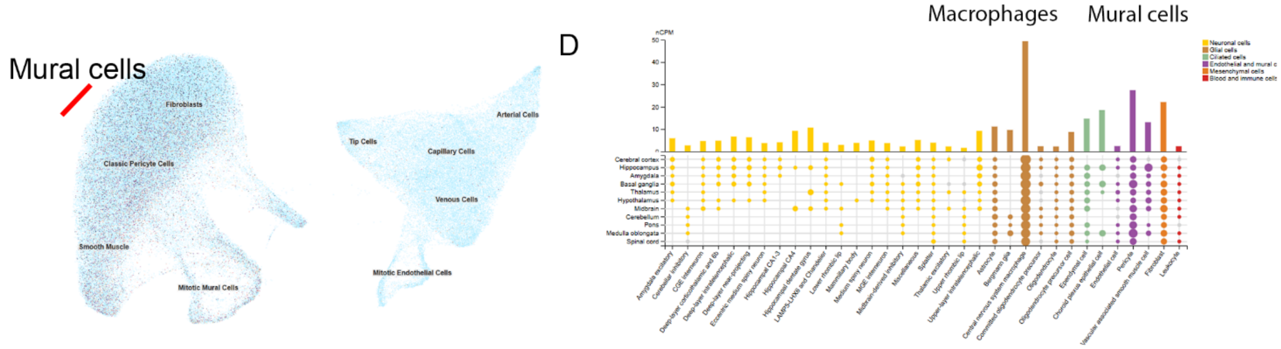


E

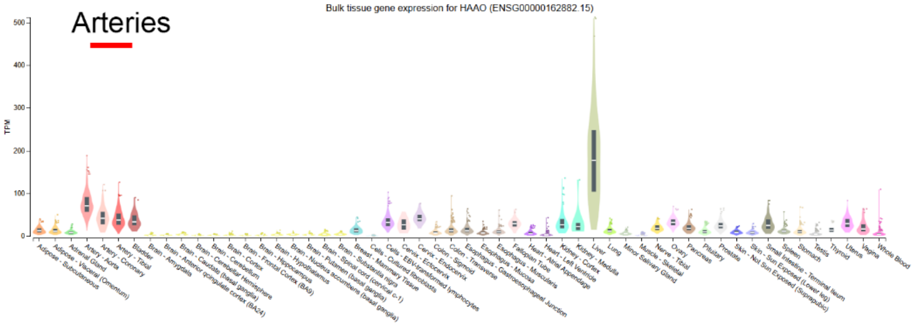


**Supplemental Fig 8: HAAO variant location**

A) Map of the HAAO gene with the variant studied. Rs1679460 is located in a Candidate Cis-Regulatory Elements (cCREs) from ENCODE (yellow bar), and in a region of H3K27Ac signal. B) Table showing the log fold change and adjusted p value as determined by the MRPA assay for this significant allele-specific enhancer. C) Expression of HAAO in human developing brain vascular mural cells[4]. F) Expression of HAAO in macrophages and mural cells in human single nucleus adult brain sequencing[5]. F) Expression in arteries and liver in the GTex dataset[6].

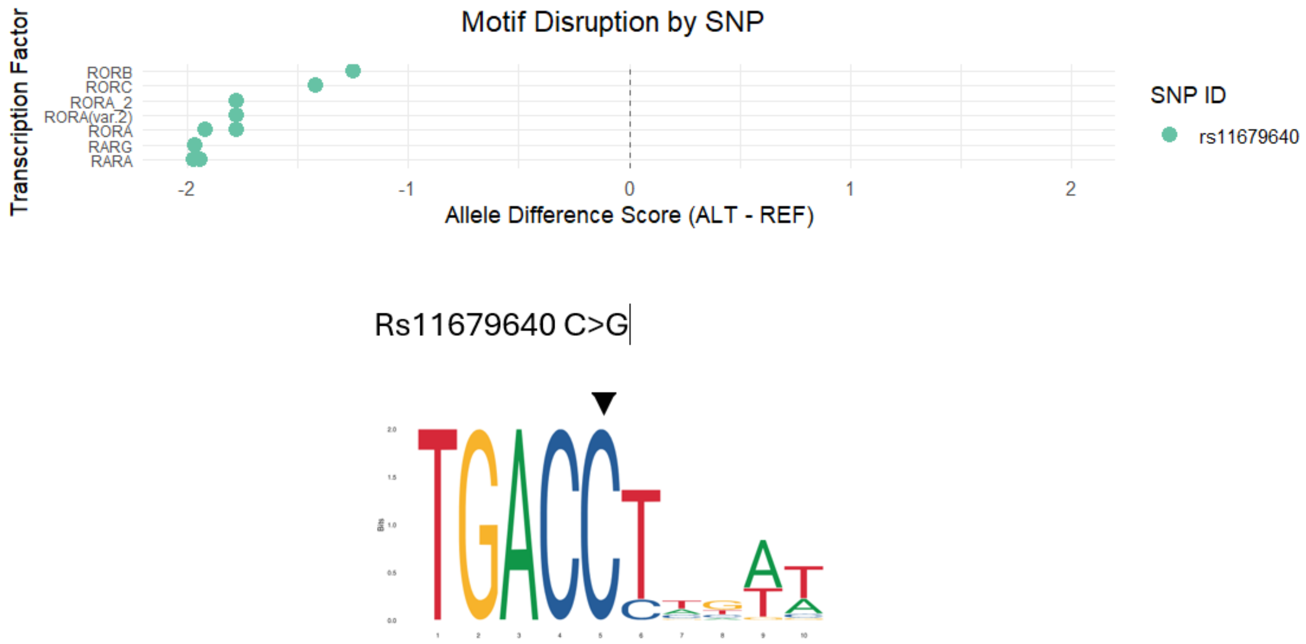


**Supplemental Fig 9: Predicted effect of SNV on transcription factor binding from MotifBreakR[7] for the HAAO rs11679640 SNV, with predicted disruptions to retinoic acid receptor (ROR, RAR) binding.** The SNV changes C>G in the highly conserved binding site for RAR Related Orphan Receptor A (RORA) from JASPAR (marked by the black arrow).

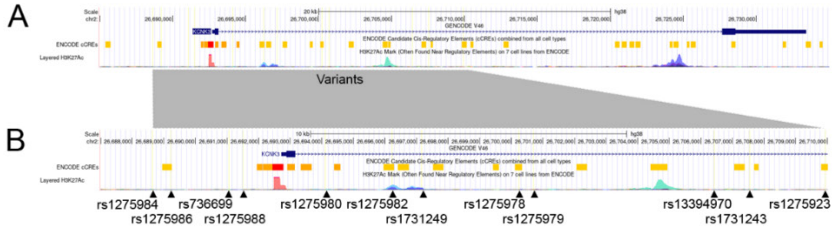


C

| SNP | logFC | adj.P.Val | Allele-specific enhancer |
|---|---|---|---|
| rs1275985 | 1.30 | 0.15703 | Not significant |
| rs35021474 | 0.33 | 0.44406 | Not significant |
| rs2586886 | 0.31 | 0.34605 | Not significant |
| rs1275978 | 3.10 | 0.00028 | Significant |
| rs1275980 | -0.66 | 0.02796 | Significant |
| rs1275986 | -0.70 | 0.03845 | Significant |
| rs736699 | 1.71 | 0.03331 | Significant |
| rs1275988 | 0.85 | 0.00000 | Significant |
| rs1275982 | 0.73 | 0.00002 | Significant |
| rs13394970 | 0.88 | 0.00005 | Significant |
| rs1731249 | 0.66 | 0.00011 | Significant |
| rs1275979 | 1.16 | 0.00023 | Significant |
| rs1275923 | 0.65 | 0.00050 | Significant |
| rs1275984 | -0.23 | 0.00286 | Significant |
| rs1731243 | 0.39 | 0.01109 | Significant |

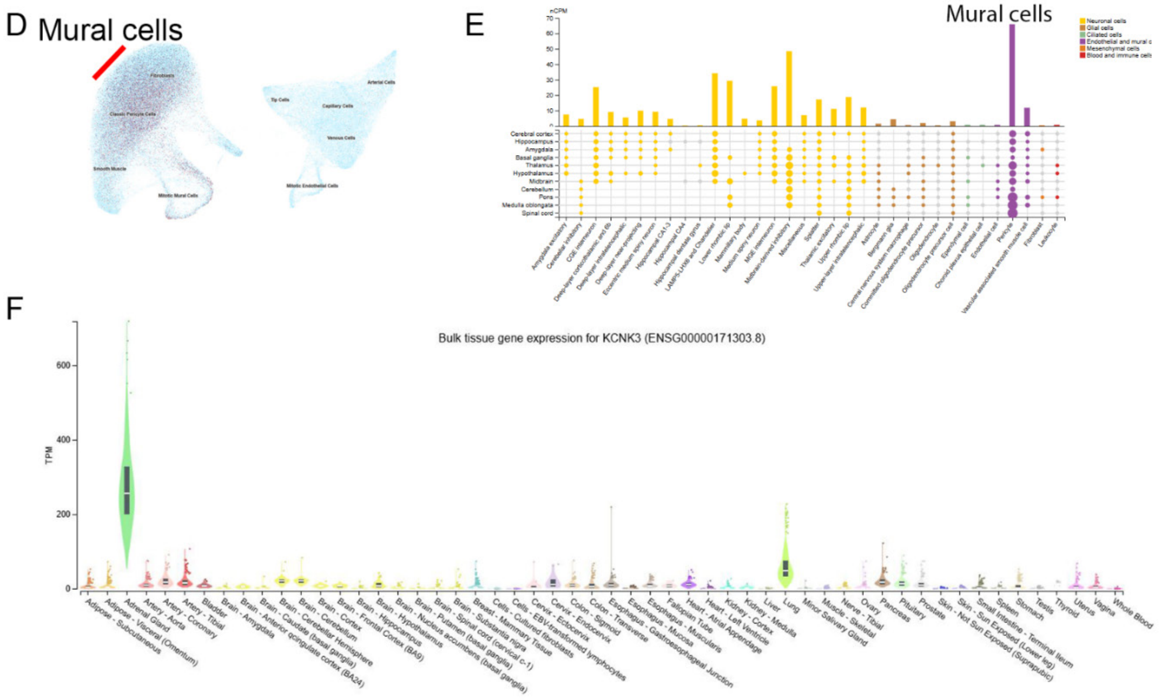


**Supplemental Fig 10: KCNK3 variant locations**

A) Map of the KCNK3 with enlargement of the region of the variants identified in our study. The grey bar shows how variants map on to the upstream and downstream intergenic regions, intron 1, and the 3’UTR of the gene. B) Some variants are located in Candidate Cis-Regulatory Elements (cCREs) from ENCODE (red and yellow bars, the variant is indicated below). C) Table showing the log fold change and adjusted p value as determined by the MRPA assay for significant allele-specific enhancers. D) Expression of KCNK3 in human developing brain vascular mural cells[4]. E) Expression of KCNK3 in neurons and mural cells in human single nucleus adult brain sequencing[5]. F) Expression in adrenal gland and lung in the GTex dataset[6].

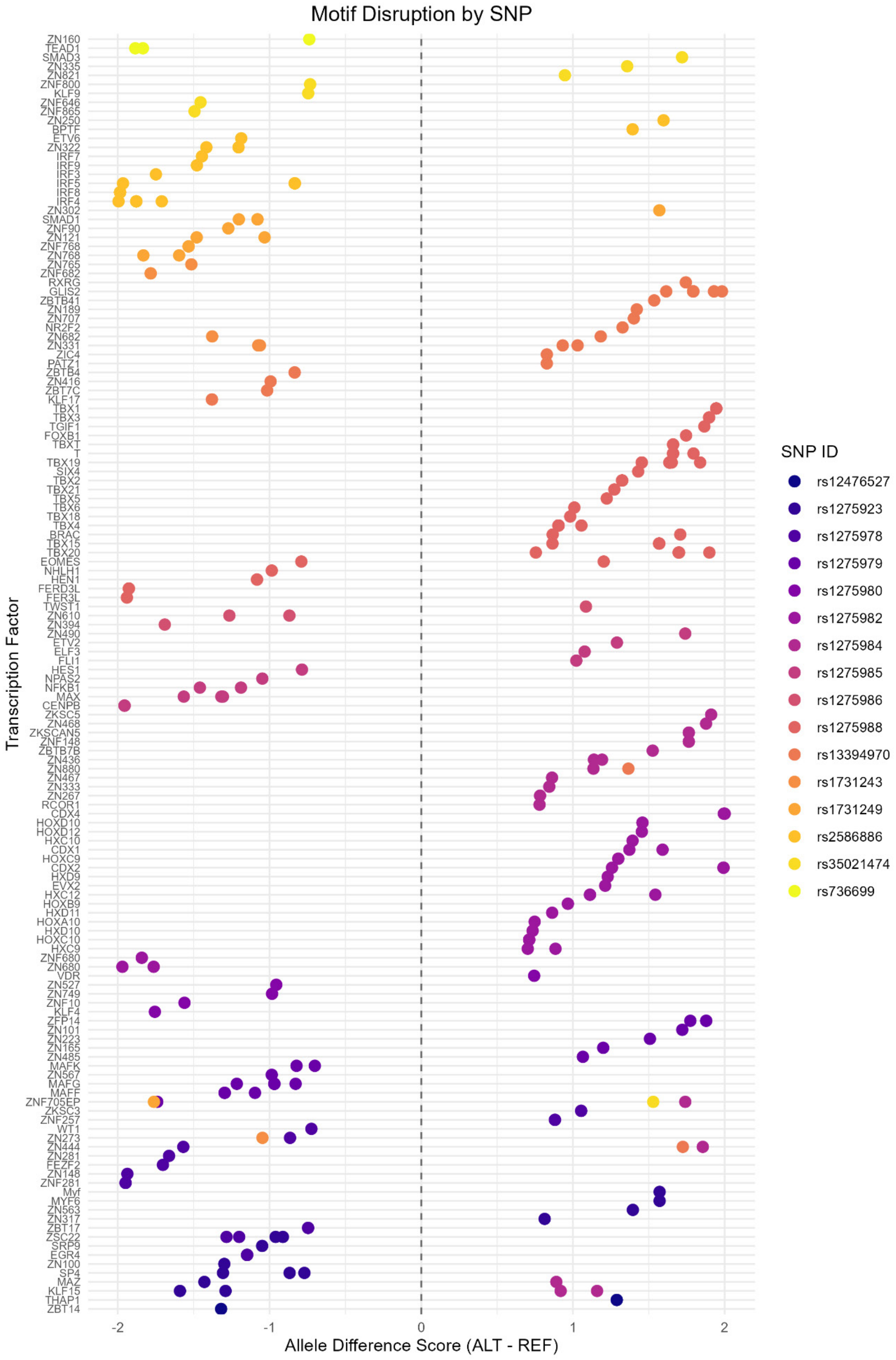


**Supplemental Fig 11: Predicted effect of SNV on transcription factor binding from MotifBreakR[7] for KCNK3 SNVs**

A

B

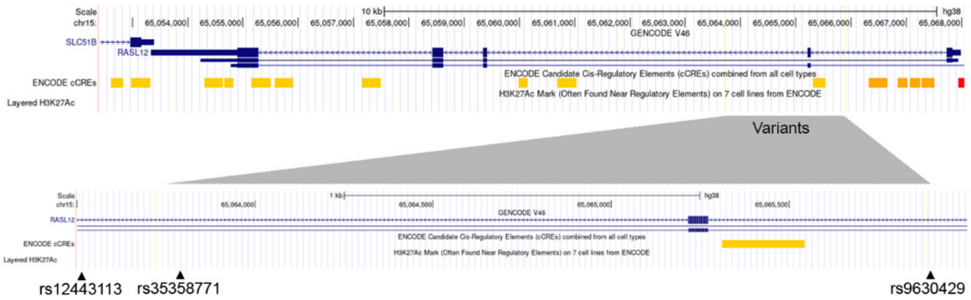


C

| SNP | logFC | adj.P.Val | Allele-specific enhancer gene | |
|---|---|---|---|---|
| rs12443113 | -0.21 | 0.46322 | Not significant | rasl12 |
| rs9630429 | 1.35 | 0.00000 | Significant | rasl12 |
| rs35358771 | 1.01 | 0.00003 | Significant | rasl12 |

D

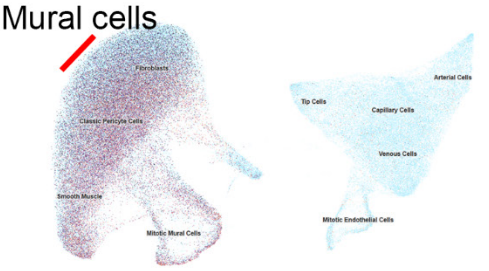


E

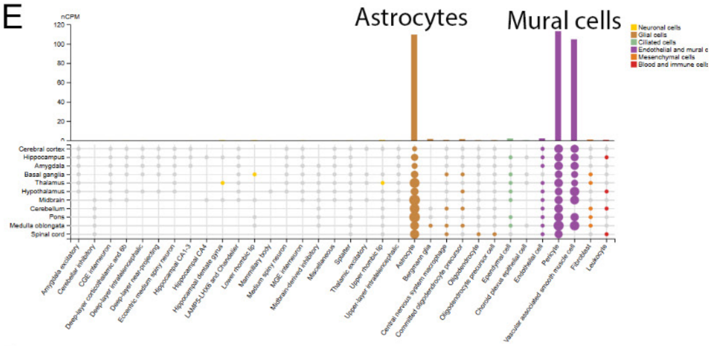


F

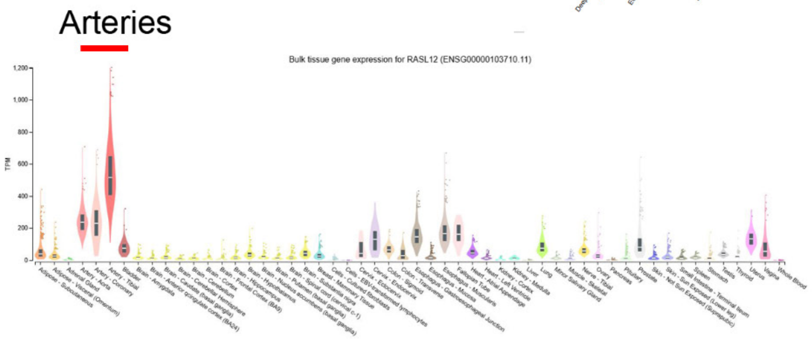


**Supplemental Fig 12: RASL12 variant locations**

A) Map of the RASL12 highlighting the region of the variants identified in our study. The grey bar shows how variants map on intron 1, and intron 2. B) None of the variants map in Candidate Cis-Regulatory Elements (cCREs) from ENCODE (red and yellow bars). C) Table showing the log fold change as determined by the MRPA assay for significant allele-specific enhancers. D) Expression of RASL12 in vascular mural cells in 1st trimester human[4]. E) Expression of RASL12 in human adult brain astrocytes and mural cells by single cell sequencing from[5]. F) Expression of RASL12 in human adult tissues from Gtex[6].

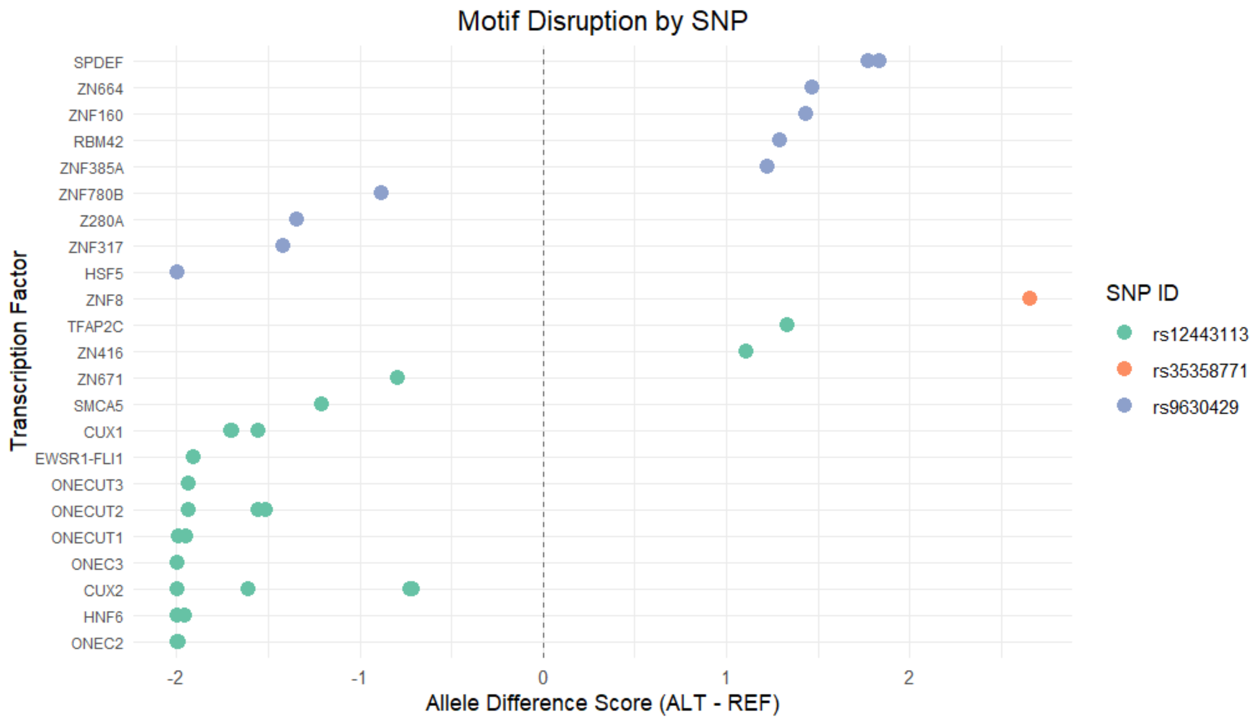


**Supplemental Fig 13: Predicted effect of SNV on transcription factor binding from MotifBreakR [7] for RASL12 SNVs**

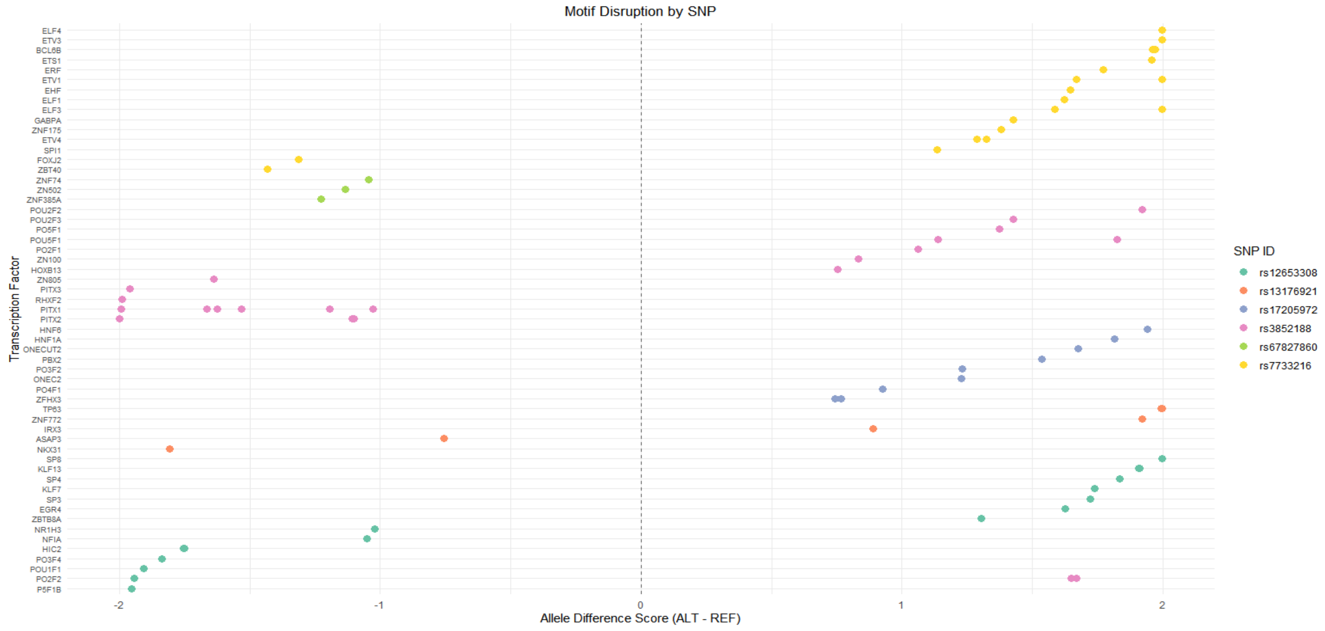


**Supplemental Fig 14: Predicted effect of SNV on transcription factor binding from MotifBreakR [7] for VCAN SNVs**

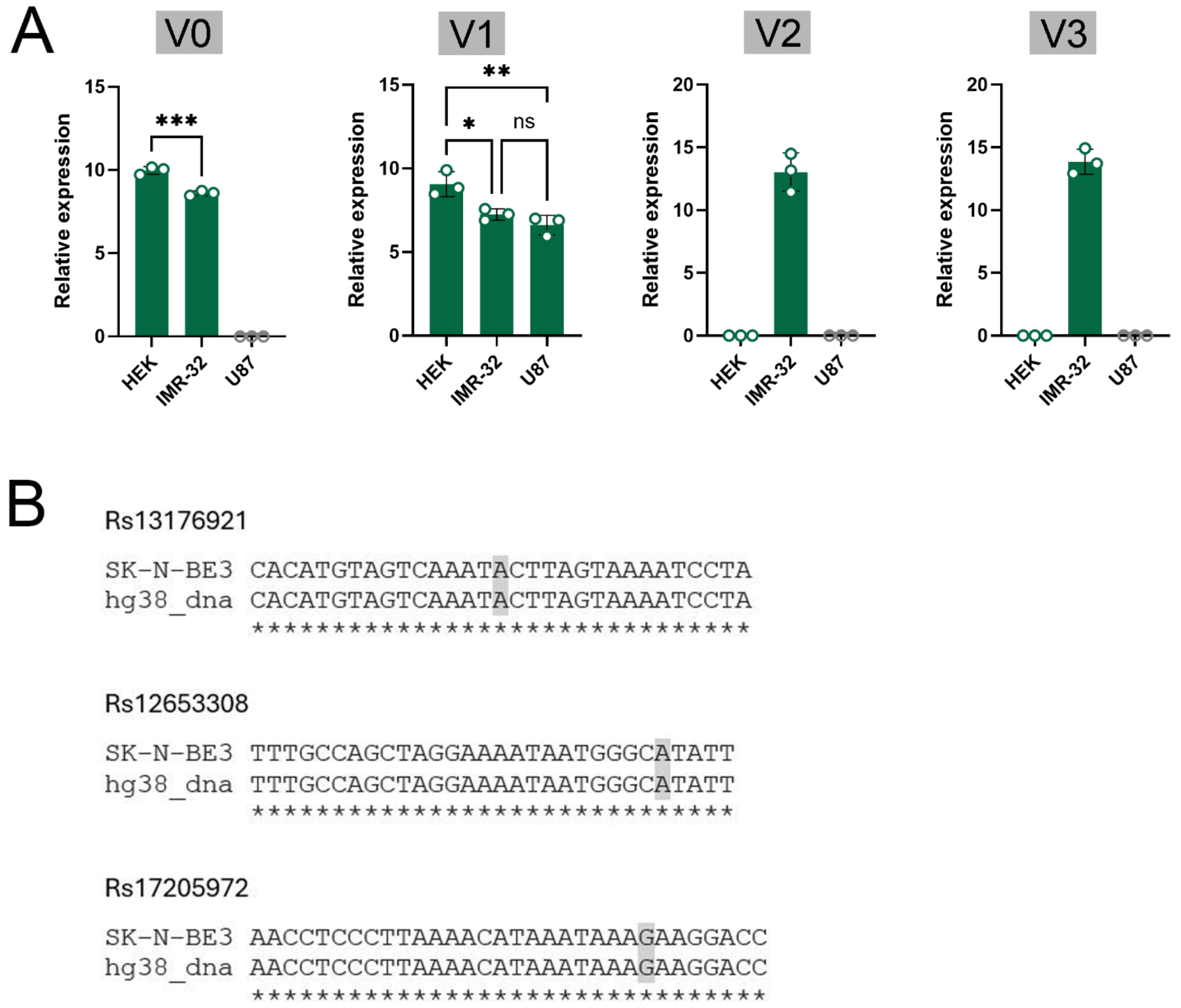


**Supplemental Figure 15: VCAN splice variant expression in cell lines identifies SK-N-BE3 cells as expressing all VCAN splice variants.**

A) Expression of VCAN variants V0- V3 in three human cell lines as determined by qPCR. N=3 biological replicates, with 3 technical replicates. Neuroblastoma line SK-N-BE3 shows expression of all 4 variants. Statistics used a One Way ANOVA with Dunnett's test. B) Sequence validation of the presence of the major allele for all 3 indicated SNVs in SK-N-BE3 cells (grey coloring).
